# TgLaforin, a glucan phosphatase, reveals the dynamic role of storage polysaccharides in *Toxoplasma gondii* tachyzoites and bradyzoites

**DOI:** 10.1101/2023.09.29.560185

**Authors:** Robert D. Murphy, Cortni A. Troublefield, Joy S. Miracle, Lyndsay E.A. Young, Aashutosh Tripathi, Corey O. Brizzee, Animesh Dhara, Abhijit Patwardhan, Ramon C. Sun, Craig W. Vander Kooi, Matthew S. Gentry, Anthony P. Sinai

## Abstract

The asexual stages of *Toxoplasma gondii* are defined by the rapidly growing tachyzoite during the acute infection and by the slow growing bradyzoite housed within tissue cysts during the chronic infection. These stages represent unique physiological states, each with distinct glucans reflecting differing metabolic needs. A defining feature of *T. gondii* bradyzoites is the presence of insoluble storage glucans known as amylopectin granules (AGs), the function of which remains largely unexplored during the chronic infection. The presence of storage glucans has more recently been established in tachyzoites, a finding corroborated by specific labeling with the anti-glycogen antibody IV58B6. The *T. gondii* genome encodes activities needed for glucan turnover inlcuding: a glucan phosphatase (TgLaforin; TGME49_205290) and a glucan kinase (TgGWD; TGME49_214260) that catalyze a cycle of reversible glucan phosphorylation required for glucan degradation by amylases. Disruption of TgLaforin in tachyzoites had no impact on growth under nutrient-replete conditions. Growth of TgLaforin-KO tachyzoites was however severely stunted when starved of glutamine despite being glucose replete. Loss of TgLaforin attenuated acute virulence in mice and was accompanied by a lower tissue cyst burden, without a direct impact on tissue cyst size. Quantification of relative AG levels using AmyloQuant, an imaging based application, revealed the starch-excess phenotype associated with the loss of TgLaforin is heterogeneous and linked to an emerging AG cycle in bradyzoites. Excessive AG accumulation TgLaforin-KO bradyzoites promoted intra-cyst bradyzoite death implicating reversible glucan phosphorylation as a legitimate target for the development of new drugs against chronic *T. gondii* infections.

**Importance:** Storage of glucose is associated with a projected need for future metabolic potential. Accumulation of glucose in insoluble amylopectin granules (AG) is associated with encysted forms of *Toxoplasma gondii*. AG which are not observed in rapidly growing tachyzoites do appear to possess glycogen, a soluble storage glucan. Here we address the role of reversible glucan phosphorylation by targeting TgLaforin, a glucan phosphatase and key component of reversible glucan phosphorylation controlling AG and glycogen turnover. Loss of TgLaforin fundamentally alters tachyzoite metabolism making them dependent on glutamine. These changes directly impact acute virulence resulting in lowering tissue cyst yields. The effects of the loss of TgLaforin on AG levels in encysted bradyzoites is heterogenous, manifesting non-uniformly with the progression of the chronic infection. With the loss of TgLaforin culminating with the death of encysted bradyzoites, AG metabolism presents a potential target for therapeutic intervention, the need for which is acute.

## INTRODUCTION

*Toxoplasma gondii* is an opportunistic protozoan parasite of all warm-blooded animals that infects one-third of humans worldwide (1, 2). Humans are primarily infected through the consumption of an encysted form of the parasite: either the oocysts shed in cat feces or tissue cysts found in undercooked meat from a chronically infected animal (3). Encysted parasites convert into tachyzoites that rapidly divide and disseminate throughout the body of the host, defining the acute phase of infection (4). Under host immune pressure, tachyzoites convert into slow-growing bradyzoites that populate tissue cysts which are found predominantly in the central nervous system and muscle, defining the chronic phase of infection (5, 6). Tissue cysts are believed to persist for the lifetime of the host and possess the ability to reactivate into tachyzoites in the context of immunosuppression. Reactivation can result in the life-threatening symptoms of toxoplasmosis, with toxoplasmic encephalitis being the primary condition leading to mortality (7, 8). The current lack of insights into bradyzoite physiology *in vivo* precludes the basic understanding needed for the development of drugs that either can clear tissue cysts or target encysted bradyzoites so as to prevent reactivation (9).

Until recently, bradyzoites within tissue cysts were considered to be dormant, metabolically inert entities. This view was challenged by our demonstration that encysted bradyzoites replicate (10, 11). Moreover, bradyzoite physiology is both diverse and complex as viewed through the lens of mitochondrial activity (12, 13), replication status (11), and, importantly, amylopectin granule (AG) accumulation (14). Although the function of AGs in bradyzoites has not been confirmed, an understanding of the roles of polysaccharides elsewhere suggests that AGs are a source of energy and biosynthetic potential needed for persistence, replication, reactivation, and transmission (15). These assumptions remain to be tested, and thus much like bradyzoites themselves, the role of AGs in the *T. gondii* lifecycle is poorly understood. Our first insights into the potential relationships connecting AG to intermediate metabolism via mitochondrial and replication activities have been exposed using imaging-based approaches (14).

AGs are large glucans found in the cytoplasm of bradyzoites that have classically served as a morphological feature distinguishing them from tachyzoites (16–19). Toxoplasma AGs are much like plant starch in that they are water-insoluble storage polysaccharides composed of branched chains of glucose (18). Unlike plant starch, however, AGs contain no detectable amylose (unbranched chains of glucose) (18). More recently, the presence of small, punctate, cytoplasmic glucans in tachyzoites that are only visible by periodic acid-Schiff (PAS) staining have been recognized (20–22), and the presence of the glucan is dependent on the *T. gondii* starch synthase (TgSS; TGME49_222800) (23). Like animal glycogen, this tachyzoite storage polysaccharide is rapidly turned over (20), as has been observed in other protozoa (24–26), and provides glucose for glycolysis (23, 27). The observation that large, insoluble glucans do not accumulate within tachyzoites as they do within bradyzoites suggests that the tachyzoite glucan could be a distinct and labile form of stored glucose, likely glycogen-like, although its exact chemical and structural identity remains unknown.

Glucose release from starch in plants requires a cycle of direct, reversible glucan phosphorylation to solubilize the starch surface, allowing access to degradation enzymes such as amylases, branching enzymes, and a phosphorylase (28–30). The cycle begins with the addition of phosphate directly to glucose by the glucan, water-dikinase (GWD) and phospho-glucan, water dikinase (PWD) that results in the unwinding of glucose chains within starch, solubilizing the starch surface (31, 32). Glucose-releasing enzymes (amylases) then degrade starch until the glucan-bound phosphate becomes a steric hindrance, at which point a glucan phosphatase is needed to remove the phosphate and reset the cycle (33–35). *T. gondii* encodes all the activities needed for glucan degradation and reversible glucan phosphorylation including the glucan phosphatase, TgLaforin (TGME49_205290) (36), and glucan dikinase, *T. gondii* GWD (TgGWD; TGME49_214260) (27). The central role of reversible glucan phosphorylation in plants is seen in *Arabidopsis thaliana* where loss of the plant glucan phosphatase, starch-excess 4 (SEX4), results in excess starch accumulation, aberrant starch morphology, and severely stunted plant growth (37, Zeeman, 1998 #1387). Additionally, loss of the glucan phosphatase, laforin, in humans, results in hyperphosphorylated glycogen that aggregates in neurons and astrocytes causing a fatal neurodegenerative childhood dementia and epilepsy (38–40). In *T. gondii*, perturbations of several genes related to glucan metabolism also result in a variety of similar defects including aberrant glucan accumulation, rewiring of central carbon metabolism, and virulence defects in mice, highlighting the central metabolic role of glucan metabolism in *T. gondii* (20–23, 27, 41–44)

In this study, we build on our understanding of reversible glucan phosphorylation and its relevance to parasite metabolism in *T. gondii*. We have recently demonstrated that TgLaforin is the glucan phosphatase in *T. gondii*, and that TgLaforin represents a unique and viable drug target (36, 45, 46). Here, we investigate the role of TgLaforin throughout the asexual stages by knocking out TgLaforin in Type II ME49 parasites. While we expected to observe effects related to the loss of TgLaforin exclusively in bradyzoites where AGs are typically observed, these effects appeared to be connected to an emerging AG temporal cycle (14). More surprisingly, the loss of TgLaforin also resulted in phenotypic effects in tachyzoites, also in a context-specific manner. We thus established a role for TgLaforin, and by extension reversible glucan phosphorylation, across both tachyzoite and bradyzoite life stages. These findings build upon previous studies that increasingly demonstrate a central role for glucan metabolism throughout the parasite’s asexual life cycle.

## RESULTS

### T. gondii tachyzoites contain a cytoplasmic glucan with a punctate distribution

Previous studies have presented biochemical evidence for rapid glucan turnover in *T. gondii* Type I RH tachyzoites (20). Moreover, small granules that stain with periodic acid Schiff reagent (PAS) have also been noted in the cytoplasm of tachyzoites (20, 21, 23). Under acid-stress conditions, these tachyzoite glucans have been biochemically characterized as pure amylopectin, and resemble AGs seen in bradyzoites (18). To further characterize the nature of this tachyzoite glucan, we used multiple methods to visualize them under unstressed, normal growth conditions (**Figure 1A**). PAS staining confirmed that Type II ME49 tachyzoites contain small punctate granules distributed throughout the cytoplasm. To determine if these PAS-stained granules in unstressed tachyzoites were more glycogen- or starch-like, they were stained with IV58B6. IV58B6 is an anti-glycogen IgM monoclonal antibody that has previously been demonstrated to be specific to glycogen (47–49) by recognizing the highly frequent branch-points found in glycogen (48). Moreover, IV58B6 does not detect other glucans such as amylopectin or amylose (the primary constituents of plant starch) (50). Tachyzoites stained with IV58B6 in a similar pattern to PAS-stained parasites, containing small punctate granules distributed throughout the cytoplasm (**Figure 1A**), suggesting that the glucan found in tachyzoites is more glycogen-like than starch-like. Finally, as is well-known, *T. gondii* tachyzoites contain almost no visible glucan within their cytoplasm when visualized by transmission electron microscopy (TEM) (**Figure 1A**), suggesting that the glucan detected by both PAS staining and IV58B6 is either water-soluble or too small to be visualized, consistent with this glucan being glycogen-like.

**Figure 1.**
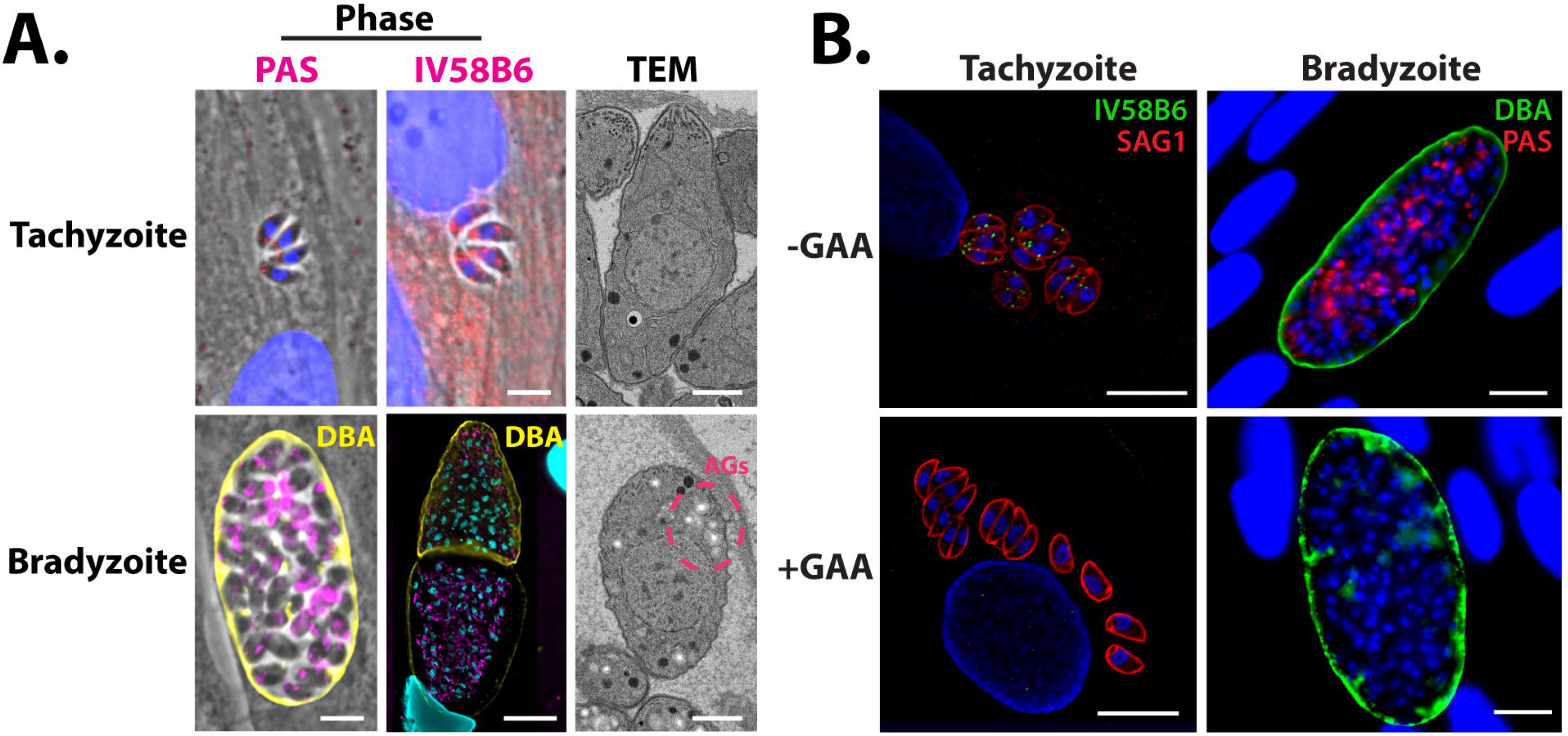
Glucan dynamics in *T. gondii* ME49 tachyzoites and in vitro bradyzoites. **A**, Microscopy-based glucan evaluation of *T. gondii* tachyzoites and bradyzoites using PAS (left), IV58B6 (middle; α-glycogen IgM mAb), and TEM (right). **B**, GAA digest of AGs in tachyzoites and bradyzoites confirms specificity of IV58B6 antibody and PAS staining. All scale bars = 5 μm.

In contrast, bradyzoites have been extensively characterized as containing starch-like AGs (17–19, 51). After *in vitro* bradyzoite conversion, much of the cytoplasm stained heavily with PAS (**Figure 1A**). Interestingly, IV58B6 staining intensity appeared to correlate negatively with *Dolichos biflorus* agglutinin (DBA) staining intensity that defines the cyst wall, implying that IV58B6 does not stain the PAS-stained glucan in bradyzoites, further reinforcing the observation that structurally distinct polysaccharides exist in tachyzoites and bradyzoites (**Figure 1A**). Finally, unlike in tachyzoites, AGs were readily identified as electron-lucent structures throughout the cytoplasm of in vitro generated bradyzoites by TEM (**Figure 1A**).

To verify the specificity of PAS and IV58B6 for glucose polymers, tachyzoites and bradyzoites were treated with acid-α-amyloglucosidase (GAA) after parasite fixation and before staining. GAA cleaves both α-1,4- and α-1,6-glycosidic bonds and can therefore completely digest glucans into glucose monomers. Indeed, GAA treatment resulted in the disappearance of staining within both tachyzoites and in vitro bradyzoites (**Figure 1B**) demonstrating their specificity for glucose polymers.

### TgLaforin colocalizes with the tachyzoite glucan

Because *T. gondii* encodes TgLaforin, a glucan phosphatase that is more animal-like than plant-like (45, 46), we reasoned that TgLaforin could be involved in the metabolism of the glycogen-like glucan found in tachyzoites. To determine if TgLaforin co-localizes with the tachyzoite glucan, endogenous TgLaforin was epitope-tagged with hemagglutinin (HA) in *T. gondii* Type II ME49ΔHXGPRT parasites (52) with a CRISPR/Cas9 mediated strategy (**Figure 2A**) (53). Successful tagging of TgLaforin was confirmed by western blotting (**Figure 2B**). Immunofluorescence analysis (IFA) of *T. gondii* tachyzoites indicated that TgLaforin is present in small puncta throughout the cytoplasm, similar to the distribution of the tachyzoite glucan (**Figure 2C**). Surprisingly, TgLaforin was not detected in *in vitro* bradyzoites by IFA, 6 days post conversion (**Figure 2C**). To verify that TgLaforin levels decrease during the tachyzoite to bradyzoite transition, we converted *T. gondii* tachyzoites to bradyzoites in cell culture using alkaline stress for 6 days and then probed the converted parasites using western blot analysis. As observed using IFA, TgLaforin-HA expression decreased dramatically over the course of bradyzoite differentiation (**Figure 2D**). Transcriptomic data from a previous study obtained from ToxoDB.org indicates that the transcript levels for TgLaforin do not substantially change over the course of differentiation, suggesting the possibility that levels of TgLaforin protein are regulated by post-translational mechanisms (54). To determine if TgLaforin colocalizes with the glucan present in tachyzoites, we co-stained TgLaforin-HA tachyzoites with either PAS or IV58B6 along with an anti-HA antibody. In tachyzoites, TgLaforin colocalized with both PAS (**Figure 2E**) and with IV58B6 (**Figure 2F**), suggesting its involvement in the metabolism of the tachyzoite glucan.

**Figure 2.**
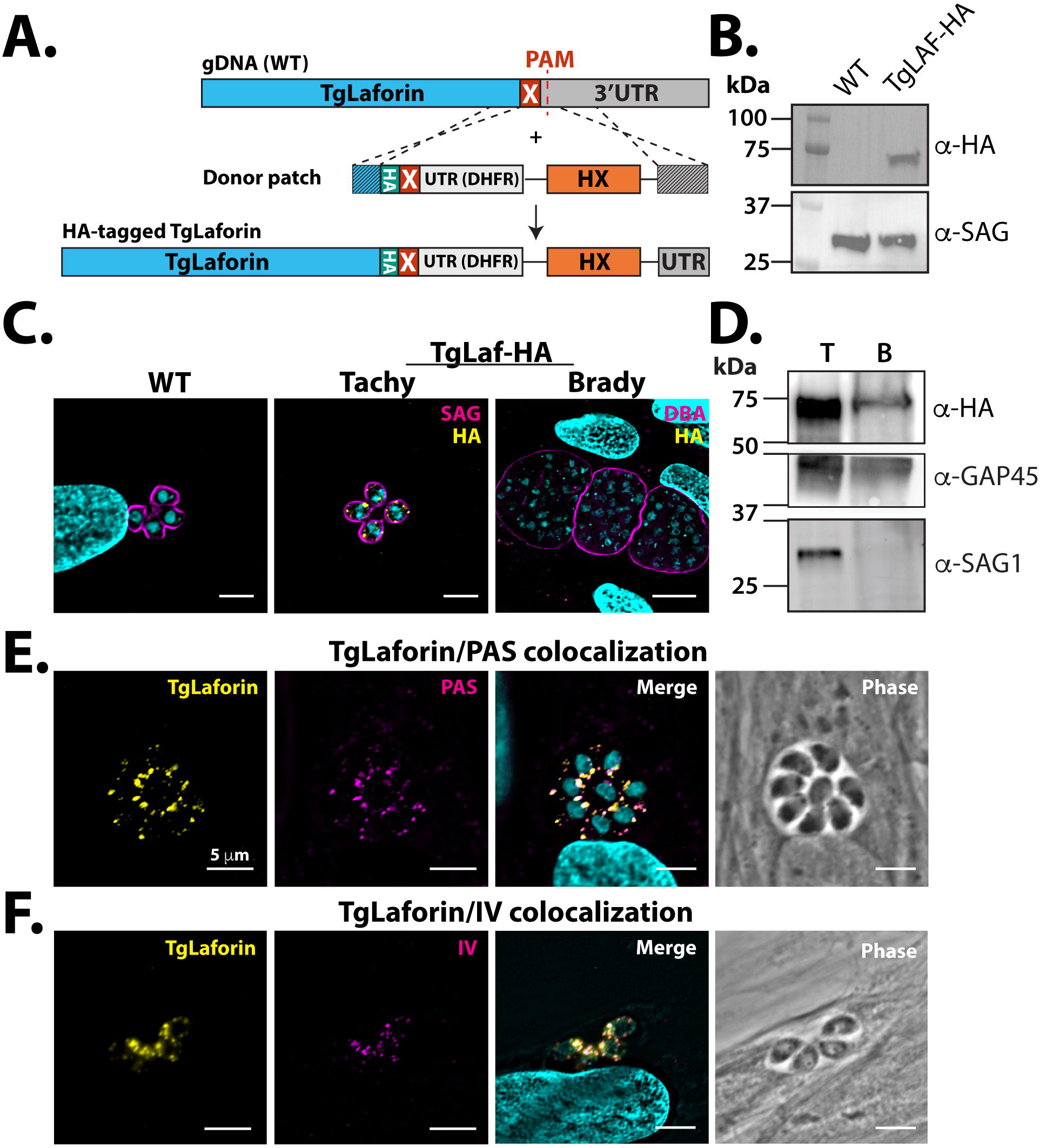
Endogenous epitope tagging and localization of TgLaforin. **A**, Schematic depicting the TgLaforin 3xHA-epitope tagging strategy. X = stop codon. **B**, Successful tagging of TgLaforin (62 kDa) was verified using immunoblot analysis with an α-HA antibody with SAG1 used as a loading control. **C**, IFA of Wild type (WT) and TgLaforin-3XHA (TgLaf-HA) tagged parasites with α-HA antibody under tachyzoite and in vitro bradyzoite conditions. Stage conversion is confirmed by Dolichos lectin (DBA) staining. **D**, Western blot analysis of TgLaforin expression levels in tachyzoites (T) and in vitro bradyzoites (B). Expression of TgLaf-HA is absent under in vitro bradyzoite conditions. GAP45 is the loading control. Decrease in SAG1 alongside increase in SRS9 confirms tachyzoite to bradyzoite conversion. **E**, IFA of TgLaforin-HA colocalization with PAS. Pearson’s coefficient: 0.765. **F**, IFA of TgLaforin-HA colocalization with IV58B6. Pearson’s coefficient: 0.737. All scale bars = 5 μm.

### Initial characterization of TgLaforin-KO tachyzoites

To dissect the role of TgLaforin in *T. gondii* glucan metabolism, TgLaforin was knocked out using CRISPR/Cas9 to disrupt the gene with a pyrimethamine-resistant form of the dihydrofolate reductase (DHFR-TS*) under a *Neospora caninum* GRA7 (NcGRA7) promoter (55) (56) (**Figure 3A**). In agreement with a genome-wide CRISPR KO screen (57), TgLaforin is a non-essential gene under standard cell culture conditions, as multiple TgLaforin-KO clones were successfully recovered. Integration of the DHFR-TS* construct into the TgLaforin locus was verified using inside/out PCR at the chimeric locus and by verifying the loss of TgLaforin transcription (**Figures 3B,C**). The TgLaforin-KO line further used in this study (designated “ΔTgLaf”) was complemented by the introduction of an epitope tagged (HA) gene driven by the TgLaforin promoter. The complementation construct was introduced at an ectopic site in the genome that lacks known coding sequences or regulatory elements on chromosome VI (**Figure S1A**), while leaving the ΔTgLaf/DHFR-TS* KO lesion intact for true complementation (58). This complemented strain, henceforth designated “COMP,” was successfully isolated and confirmed by PCR (**Figure S1B**). Expression levels and localization were similar to those seen in the TgLaforin-HA line as confirmed by western blotting and IFA (**Figures S1C-D**),

**Figure 3.**
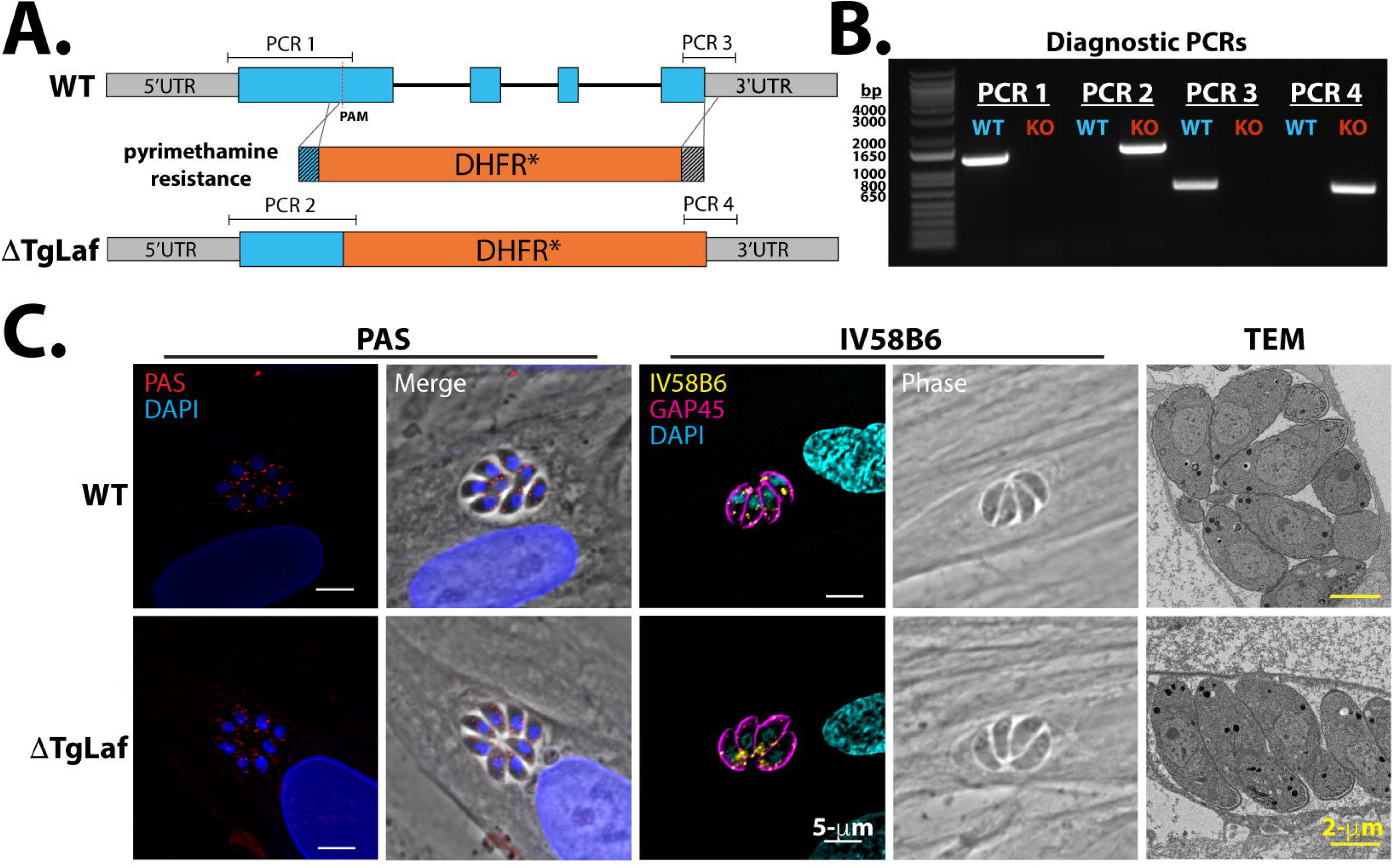
Loss of TgLaforin results in no gross glucan abnormalities under glucose replete conditions. **A**, Schematic of TgLaforin KO strategy: the pyrimethamine-resistance DHFR* gene containing 40-nt homologous arms (dark gray boxes on either end of gene) was inserted into the TgLaforin locus via homologous recombination. A double stranded break was induced using CRISPR/Cas9-GFP with a PAM site in the first exon. **B**, Inside/out PCR verification of DHFR integration into the TgLaforin locus. Amplicons (PCR1-4) are illustrated in (A). **C**, Loss of TgLaforin mRNA was confirmed by amplifying full-length TgLaforin cDNA generated from both WT and ΔTgLaf strains. Actin cDNA amplification serves both as a loading control and as a control to verify the absence of gDNA. **D**, Analysis of glucan levels in ΔTgLaf tachyzoites using three different approaches: PAS and IV58B6 immunofluorescence staining, and TEM.

To evaluate effects of a TgLaforin-KO, glucan levels in WT and ΔTgLaf tachyzoites were first compared using our suite of glucan detection techniques (**Figure 3D**). Surprisingly, the size and number of PAS-stained granules were not significantly changed in ΔTgLaf tachyzoites relative to WT parasites. Levels of IV58B6 also remained unaltered after the loss of TgLaforin, and no aberrant glucan accumulation was observed by TEM as has been previously reported when genes related to AG or central carbon metabolism were knocked out in *T. gondii* tachyzoites (20–23, 41, 43, 44) (**Figure 3D**)

Loss of glucan phosphatases in plants and animals results in aberrant glucan accumulation, and such a phenotype was not observed within tachyzoites under standard growth conditions.

### Loss of TgLaforin results in upregulation of glutaminolysis and glutamine dependence in tachyzoites

Glucan catabolism is significantly affected by the presence of covalently bound phosphate, and, therefore, loss of glucan phosphatases has profound downstream metabolic impacts in other systems (59, 60). We thus speculated that loss of TgLaforin would result in the reduced efficiency of glucan utilization in tachyzoites and also affect downstream central carbon metabolism. To test this hypothesis, we used gas chromatography/mass spectrometry (GC/MS) steady-state metabolomic analysis of 3 μm filter-purified, syringed-passaged intracellular tachyzoites employing a previously developed sample preparation technique (61).

Previously, it was demonstrated that *T. gondii* tachyzoites primarily utilize glucose and glutamine to drive central carbon metabolism, synthesize macromolecules, and proceed normally through the lytic cycle (62). Glucose primarily fuels glycolysis, and glutamine undergoes glutaminolysis to drive the tricarboxylic acid (TCA) cycle. In the absence of glucose, *T. gondii* can upregulate both glutaminolysis and gluconeogenesis to make up for the loss of glucose (62, 63).

While ΔTgLaf metabolite levels remained unaltered relative to WT tachyzoites across much of the TCA cycle, steady-state levels of metabolites immediately downstream of glutamine were consistently more abundant in ΔTgLaf parasites compared to their WT counterparts (**Figure 4A**), supporting our hypothesis that ΔTgLaf parasites were deficient in glucan/glucose utilization. An increase in metabolites downstream of glutamine in ΔTgLaf parasites demonstrates that ΔTgLaf parasites are possibly compensating for deficiencies in glucose metabolism, supporting a role for the tachyzoite glucan in intermediate *T. gondii* glucose metabolism.

**Figure 4.**
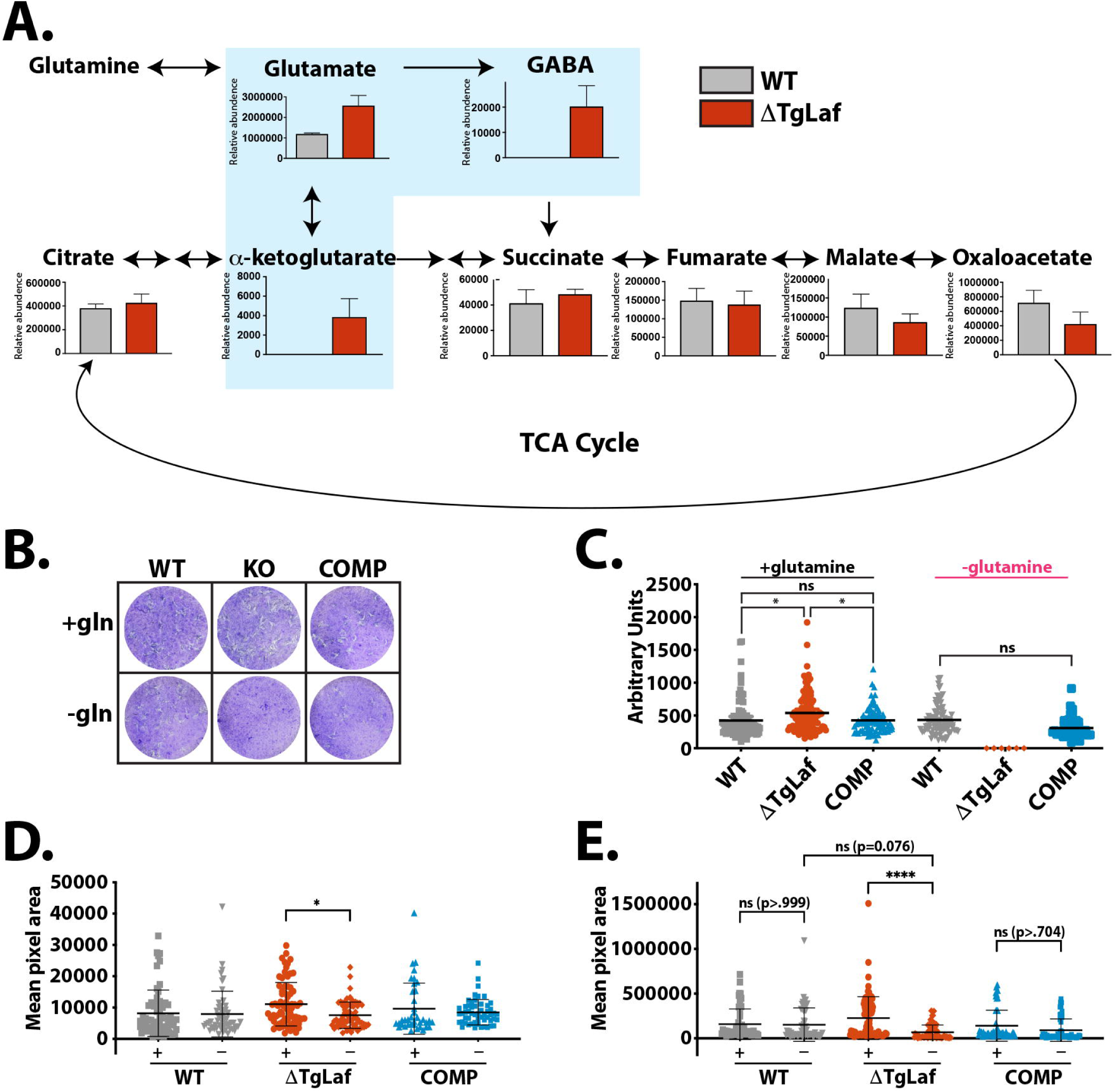
ΔTgLaf parasites are dependent on glutamine for normal plaque formation. **A**, Steady-state metabolomics suggests upregulation of glutaminolysis in ΔTgLaf parasites. Metabolite levels of intracellular tachyzoites were analyzed after 48 hours of growth in HFFs by GC/MS analysis. Data were collected from 3 independent replicates. Statistical comparisons were done using unpaired two-tailed t-tests. Statistical significance is as follows: **p<0.01, ns=p>0.05, nd=not detected. **B**, Representative plaque assays following staining of infected HFF monolayers with crystal violet under nutrient replete (+gln) and glutamine free (gln-) conditions. WT, ΔTgLaf (KO) and complemented (COMP) lines were evaluated. **C**, Quantification of visible plaques for WT, ΔTgLaf and COMP lines under nutrient replete and glutamine free conditions represented as the plaque area (pixels) was measured across three independent replicates 6 days following infection. **D**, Pixel area of nascent plaques/vacuoles after 3 days of growth as monitored by IFA microscopy following staining with the PVM maker GRA3 under both glutamine replete (+) and depleted conditions (-). **E**, Pixel area of plaques monitored as in (D) after 6 days of growth. Statistical comparisons for C-E were done using an ordinary one-way ANOVA using Tukey’s post-hoc test to correct for multiple comparisons. Statistical significance is indicated as follows: *p<0.05, ****p<0.0001, ns=p>0.05.

To determine if loss of TgLaforin resulted in increased dependence on glutamine due to impaired access to glucose, we performed plaque assays in the presence and absence of glutamine (**Figure 4B**). In replete media, ΔTgLaf parasites established a similar number of plaques (data not shown), indicating no defect in infectivity. ΔTgLaf plaques were slightly larger than both the WT and COMP lines after 10 days of growth (**Figure 4C**). To test the effects of glutamine starvation on ΔTgLaf parasites, glutamine was removed from plaque assays after parasite invasion to evaluate the effects of glutamine removal on parasite growth independent of the initial invasion event. In the absence of glutamine, ΔTgLaf parasites were unable to form visible plaques, whereas both the WT and COMP parasites formed plaques comparable to those formed under glutamine-replete conditions (**Figures 4B, C**).

### TgLaforin is required for repeated rounds of progression through the lytic cycle

The absence of differences in plaque number suggest that there is no specific defect in infectivity. To determine which aspects of the *T. gondii* lytic cycle were impaired in the absence of glutamine, the effects of glutamine starvation on initial parasite replication and egress (stimulated with both A23187 (64) and zaprinast (65)) were evaluated. In both assays, intracellular parasites were pre-starved of glutamine for at least 72 h before assay initiation. Surprisingly, glutamine starvation had no effect on stimulated egress or initial parasite replication across the three lines (**Figures S2A-C**). These data demonstrate that the absence of plaques under glutamine deficient conditions cannot be pinpointed to a single aspect of the ΔTgLaf lytic cycle, and that the reason for the apparent absence of plaques manifested later in the infection cycle.

Plaques develop due to repeated cycles of localized infection and cell lysis resulting in the clearance of infected cells over time. The absence of visible clearance prompted us to examine infected host-cell monolayers for clusters of infected cells using a higher magnification than is typically used in a traditional plaque assay. Low numbers of parasites were seeded onto glass coverslips and fixed at 3- and 6-days post-infection, allowing for visualization of developing plaques at a high magnification. In these experiments, glutamine-depleted host cells were pre-starved of glutamine prior to infection with parasites to allow for potential for invasion defects. Importantly, ΔTgLaf parasites demonstrated similar infectivity to WT parasites under both glutamine-replate and depleted conditions, indicting no gross initial invasion defect. After 3 days of growth, no statistical differences of nascent plaque sizes were noted between glutamine-replete and starved conditions in both the WT and COMP lines. However, ΔTgLaf parasites in glutamine starved conditions were already 1.5x smaller in area than their counterparts in replete conditions (**Figure 4D**). By day 6 of growth, this difference had widened to a >3x difference between glutamine replete/depleted ΔTgLaf parasites (**Figure 4E**). Such a difference was not detected between the two conditions in WT/COMP parasite lines. By measuring the internal clearing area relative to the total plaque perimeter, it was also noted that ΔTgLaf parasites were much less capable of forming clearings than the WT/COMP lines (**Figures S2D, E**), rather they formed clusters of infected cells akin to “turbid plaques” (66, 67) due to their presumed inability to compete with host cell growth, as the infection progressed. This observation explains the apparent absence of plaques seen at the lower magnification used in traditional plaque assays (**Figure 4B**). The modified plaque assay therefore confirmed that the loss of TgLaforin penalized the summation of repeated rounds of the energy-demanding lytic cycle rather than one particular aspect of the lytic cycle. Representative images from this assay can be found in **Figure S2D**.

### Loss of TgLaforin results in aberrant bradyzoite AGs in vitro

To determine if loss of TgLaforin resulted in bradyzoite conversion defects, or aberrant AG accumulation, parasites were converted to bradyzoites *in vitro* using alkaline stress. During differentiation, the parasitophorous vacuole membrane (PVM), delimiting the replicative niche established by tachyzoites, converts into the cyst wall that surrounds bradyzoites within their host cell (68, 69). The cyst wall is heavily glycosylated and contains N-acetylgalactosamine (Gal-NAc) that is detectible with Dolichos biflorus agglutinin (DBA) (68). Using DBA-FITC intensity as a marker for differentiation, no penalty was imposed by the loss of TgLaforin on cyst wall formation over the course of six days (**Figure 5A**). Somewhat surprisingly, ΔTgLaf mutant parasites tended to exhibit stronger labeling with DBA at day 6. We additionally assessed the levels of accumulated glucans using PAS staining (**Figure 5B**). Semi-quantitative analysis of PAS intensity within vacuoles during stage conversion showed an expected increase over time, but no significant difference between the WT and ΔTgLaf parasites was detected over the time course examined.

**Figure 5.**
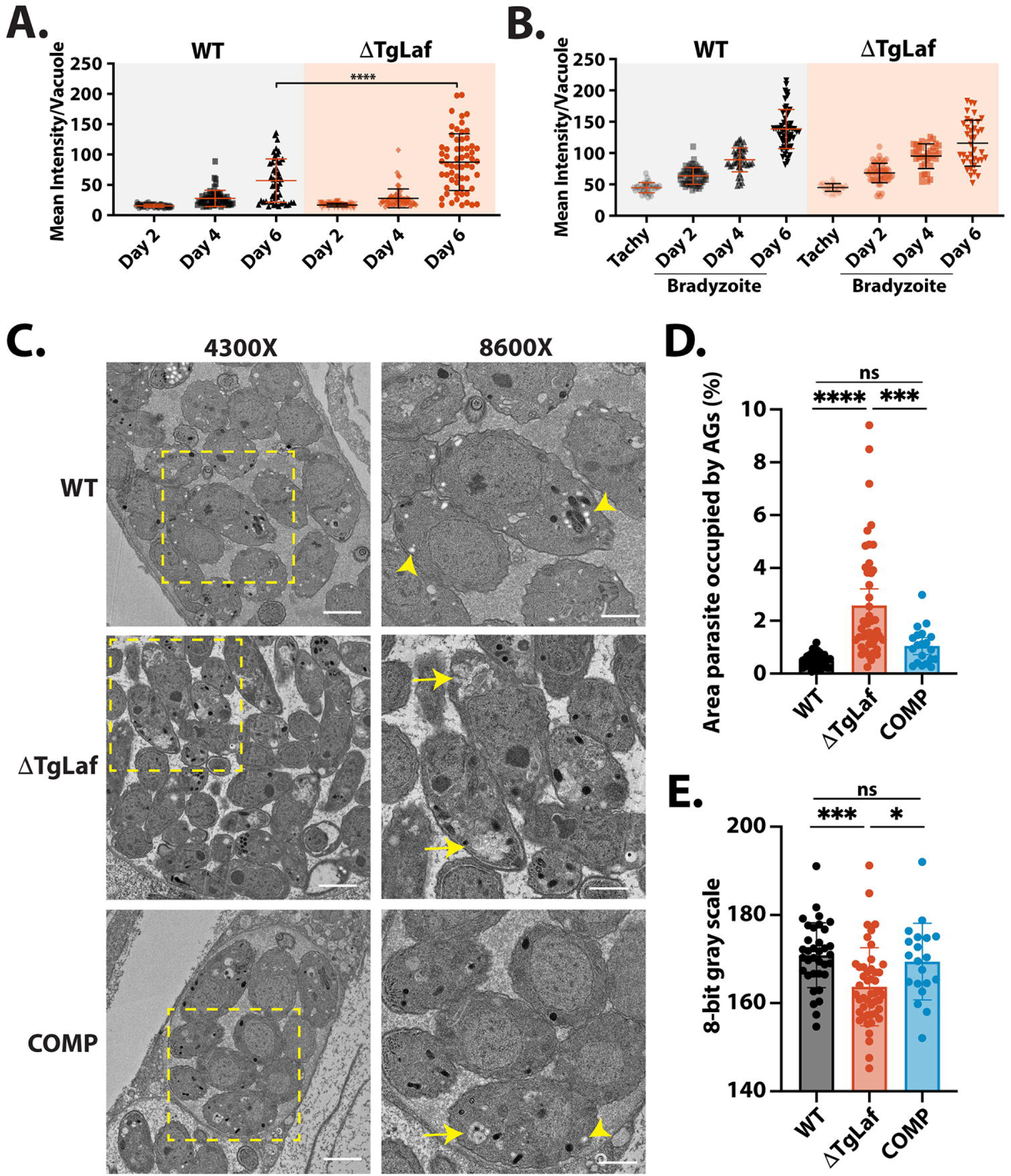
Loss of TgLaforin results in aberrant glucan morphology and accumulation in *in vitro* bradyzoites only visible by TEM. **A**, *In vitro* tachyzoite to bradyzoite conversion efficiency of ΔTgLaf vs WT parasites as measured by DBA intensity confirms no defect in induced stage conversion in ΔTgLaf parasites. **B**, Change in AG levels detected using the intensity of PAS labeling during bradyzoite conversion reveals no statistically different levels in AG accumulation based on PAS intensity. **C**, Representative TEM images of bradyzoites from each indicated parasite line. At 4300x magnification, scale bar = 2 μm; at 8600x magnification, scale bar = 1 μm. Arrowhead = canonical AG (white, round/ovoid); Arrow = aberrant AG (grey, flattened, multi-lobed). **D**, Quantification of relative parasite AG content and **E**, AG grayness across parasite lines using 8-bit grayscale (0=black, 255=white). Statistical comparisons were done using an ordinary one-way ANOVA with Tukey’s post-hoc test to correct for multiple comparisons. Statistical significance is indicated as follows: *p<0.05, ***p<0.001, ****p<0.0001, ns=p>0.05.

Because PAS is not specific to glucans and can stain other glucose-containing molecules such as glycosylated protein and provides no resolution on glucan morphology, we utilized TEM to gain higher resolution on AG formation during bradyzoite differentiation. After 6 days of conversion, WT parasites produced AGs that were circular/ovoid and white (**Figure 5C**). In contrast, ΔTgLaf parasites contained irregular AGs that were morphologically distinct from AGs that were observed in WT parasites (**Figure 5C**). AGs in ΔTgLaf parasites appeared amorphous and grayer, while appearing to occupy more area of the parasite cytoplasm compared to WT parasites. To quantify this phenotype, the area of AGs was calculated relative to total parasite area to determine the percentage of the parasite body occupied by AGs in both WT and ΔTgLaf strains (**Figure 5D**). Strikingly, AGs occupied approximately 4x more relative area in ΔTgLaf parasites when compared to WT, indicating that PAS staining may lack the sensitivity to capture this difference, in tissue culture generated bradyzoites. When analyzed on an 8-bit gray scale, AGs in ΔTgLaf parasites were significantly grayer (grayscale 0-255 is black-white) than those found in WT parasites, highlighting potential chemical differences (such as predicted hyperphosphorylation) resulted in differential interactions of ΔTgLaf AGs with the TEM contrast reagents, likely the heavy metals used in processing (**Figure 5E**).

Examination and quantification of AGs in the COMP line revealed that complementation of TgLaforin restored most of the circular/ovoid AG cross sections while they also occupied less space in the cytoplasm and were overall more like those found in WT parasites (**Figures 5C-E**). Thus, cell culture experiments demonstrate that the loss of TgLaforin presents itself in both a context and life cycle stage-specific manner.

### Loss of TgLaforin results in attenuated virulence and cyst formation in vivo

We hypothesized that loss of TgLaforin may impose a steep penalty under the stresses and potential nutrient scarcities encountered *in vivo* as it does when nutrients (such as glutamine) are scarce *in vitro*. To test this hypothesis, equal numbers of male and female CBA/J mice were infected with 100 tachyzoites intraperitoneally (i.p.) and monitored daily using a previously developed five-stage body index score to track the severity of symptoms associated with a tachyzoite infection over the course of 28 days (52).

Mice infected with WT parasites began demonstrating symptoms of infection ten days after infection with tachyzoites (**Figure 6A**). However, mice infected with ΔTgLaf parasites did not begin to exhibit symptoms until 15 days after infection. Moreover, mice that became symptomatic from WT parasite infections often proceeded through all stages of symptomology, and only a minor proportion of mice that became sick were able to recover from infection (>70% of mice became moribund or died). Infection from ΔTgLaf parasites, however, resulted in the majority of mice only developing mild symptoms (Stage 2 or less) with many of these mice recovering (**Figure 6A**). The attenuated capacity of the ΔTgLaf parasites to cause symptoms in mice was reflected in the mortality rates of the infected mice: infection with WT parasites resulted in 73% mortality rate after 28 days whereas ΔTgLaf parasites only caused 17% mortality (**Figure 6B**). Complementation of TgLaforin partially rescued this defect in virulence as COMP parasites resulted in an earlier onset of symptomatic infection at Day 11, and the majority (53%) of mice succumbed to infection during the first 28 days (**Figures 6A, B**).

**Figure 6.**
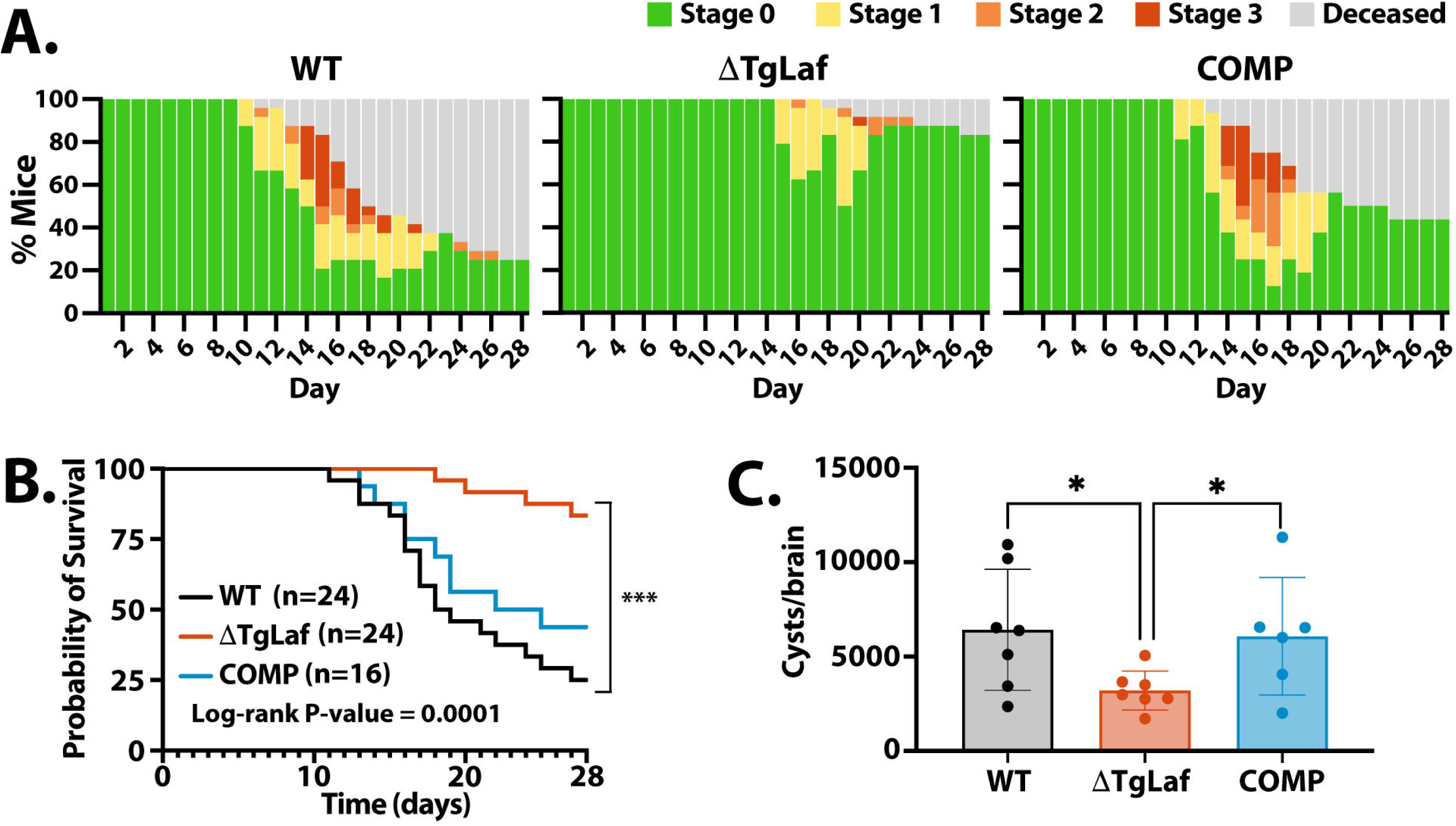
Loss of TgLaforin attenuates tachyzoite virulence and cyst burden in mice. Equal numbers of male and female CBA/J mice were infected i.p. with 100 tachyzoites from each group and monitored for both symptom progression and mortality. **A**, Symptomology throughout the acute (< day 20) and early chronic phases of infection (day 21-28). Mice were monitored 1-2x/day and assigned a body score index ranging from asymptomatic (Stage 0) to moribund/deceased. Staging of disease progression is discussed in (52). **B**, Kaplan-Meier curve of mouse survival throughout acute and early chronic infection following tachyzoite infection. **C**, A cohort of mice that survived 4- weeks were euthanized, and the cyst burden determined as done previously described (70). Error bars depict SD from the mean. Statistical comparison for Kaplan Meier curves is indicated on plot, and statistical comparison of cyst burden was done using unpaired two-tailed t-tests. Statistical significance: *p<0.05, ***p<0.0002.

Because the acute phase of infection was significantly attenuated by the loss of TgLaforin, we hypothesized that cyst numbers would be significantly lowered. To determine the number of cysts formed after 28 days of acute infection (Week 4), we used a previously established protocol for harvesting and counting tissue cysts from infected mouse brains, following purification on Percoll gradients (11, 70). Consistent with the ability of ΔTgLaf parasites to stage convert in culture, mutant parasites were able to establish tissue cysts *in vivo*. However, the number of cysts recovered from ΔTgLaf infected animals was lower than those obtained from WT infected animals (**Figure 6C**). Restoration of TgLaforin in the COMP line effectively restored tissue cyst yields.

### **Δ**TgLaf tissue cysts can reestablish infections in naïve mice

To determine if the loss of TgLaforin impacted the overall viability/infectivity of *in vivo* tissue cysts, we examined the disease progression in WT, ΔTgLaf, and COMP infected animals following injection of 20 tissue cysts i.p. Consistent with prior data (52), infection with tissue cysts results in markedly lower pathology and consequent mortality during the acute phase for WT as well as both the ΔTgLaf and COMP lines (**Figure 7A, B**). Mortality from cyst infections did not differ statistically among the three lines (**Figure 7A, B)**. Twenty-eight days post-infection, cyst burdens were again enumerated for each line. ΔTgLaf parasites were once again much less competent at forming cysts *in vivo* (**Figure 7C**). However, unlike the tachyzoite infection, the COMP line was unable to rescue this defect in cyst formation (**Figure 7C**), suggesting that physiological and metabolic changes associated with the loss of TgLaforin manifest differently based on the life cycle stage, impacting their capacity to be complemented.

**Figure 7.**
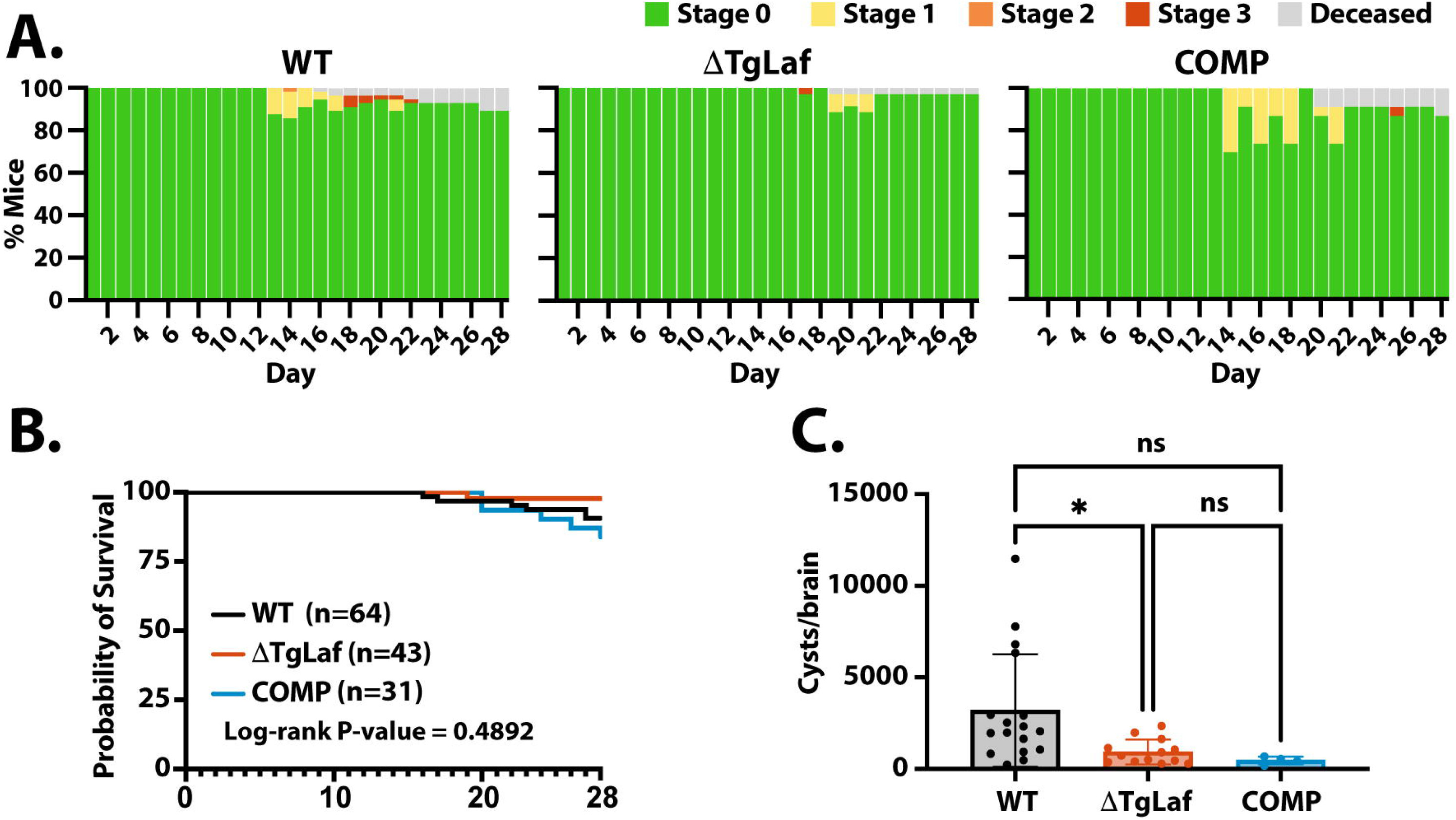
Mouse infection with ΔTgLaf tissue cysts results in milder illness and lower cyst burden. CBA/J mice were infected i.p. with 20 cysts taken from previously infected mouse brains and monitored for symptomology and death. Tissue cysts were harvested from mice infected for 4 or 6 weeks respectively. **A**, Symptomology throughout the acute and early chronic phase of infection as described in Figure 6B. **B**, Kaplan-Meier curve of mouse survival during the acute and early chronic infection reveal low overall mortality associated with cyst initiated infections for all parasite lines. **C**, Tissue cyst burden following a 4-week bradyzoite initiated infection. Cyst numbers were determined as described in Figure 6C. Statistical comparison of cyst burden was done using unpaired two-tailed t-tests. Statistical significance: *p<0.05, ns=p>0.05.

### Loss of TgLaforin results in a delayed starch-excess phenotype within encysted bradyzoites

We leveraged the inherent fluorescence of PAS coupled with an optimized labeling protocol to determine the mean intensity of WT, ΔTgLaf and complemented (COMP) tissue cyst using Image J (**Figure. 8A**). Similar to the pattern observed with in vitro generated bradyzoites (**Figure. 5**), *in vivo* derived tissue cysts harvested at 28 days (Week 4) post infection failed to demonstrate a starch-excess phenotype in the ΔTgLaf mutant (**Figure 8A**). Notable here, is the fact that at week 4 post infection, bradyzoites are in a state of low AG accumulation (14), and presumably, low turnover, potentially minimizing the impact of the loss of TgLaf. Our prior data indicate that tissue cysts harvested at week 6 post infection are in markedly more dynamic state noted by increased accumulation of AG, higher levels of active mitochondria and increased replicative activity (14). Quantification of mean PAS intensity exposed a dramatic increase in AG levels comparing WT and ΔTgLaf tissue cysts, with overall levels between WT and COMP cysts remaining not significantly different (**Figure 8A**). This points to the effect of the loss of TgLaf demonstrating a delayed starch excess phenotype, indicating the penetrance of the mutation is potentially connected to achieving a threshold level of AG as a part of an emerging AG cycle (14).

**Figure 8.**
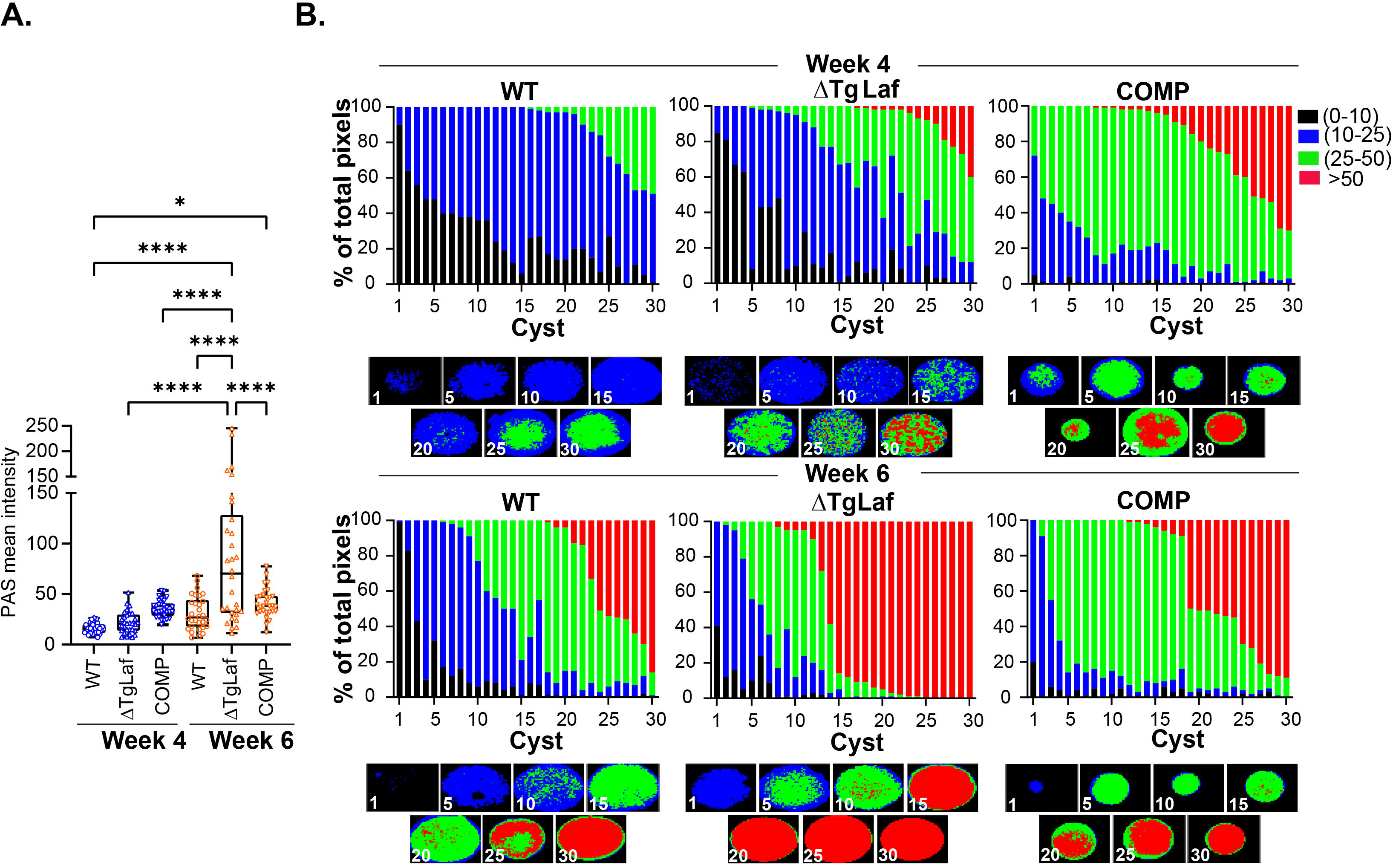
Quantification of amylopectin levels in tissue cysts based on PAS intensity. **A.** The mean intensity of PAS labeled tissue cysts measured using Image J for WT, ΔTgLaf and COMP tissue cysts harvested at 4 and 6 weeks post infection reveal no difference in AG levels across these parasite lines at week 4. Highly significant differences are evident between WT/COMP and DTgLaf tissue cysts at week 6 post infection indicating a temporal component for the emergence of the starch excess phenotype. n=30 for each parasite line at each time point of harvest. Statistical analysis was done using Tukey’s multiple comparison test. P values: * p <0.05, **** p< 0.0001 **B.** Analysis of WT, DTgLaf and COMP tissue cysts harvested at weeks 4 and 6 using AmyloQuant. The 30 tissue cysts per sample presented in A. were analyzed using AmyloQuant. While acquired at random, the data are arrayed from low to high PAS intensity presented as the percentage of pixels in each classification bin. The classification bins defined as: background: Black: 0-10 grayscale, low: Blue: 10-25 grayscale, intermediate: Green 25-50 grayscale and high intensity pixels: Red >50 grayscale. The higher overall sensitivity of AmyloQuant reveals increased accumulation of AG at week 4 in the DTgLAf and COMP lines. This phenotype is greatly exaggerated at week 6 post infection consistent with a temporal component for the phenotypic manifestation of the starch excess phenotype. The pattern in the COMP parasites is intermediate between the WT and ΔTgLaf cysts. The distribution of PAS intensity levels within the imaged tissue cysts generated by AmyloQuant for cysts in 5 cyst intervals is presented under each set of stacked plots.

The development of AmyloQuant, an imaging-based application, permits the direct measurement of relative AG levels within tissue cysts (14). With is application, individual PAS-labeled pixels are classified into 4 bins defining background pixels (black: 0-10 grayscale), low intensity (blue: 10-25), intermediate (green: 25-50) and high (red: > 50 grayscale). Thirty randomly acquired tissue cysts, (the mean intensity of which were reported in **Figure 8A**) from WT, ΔTgLaf and COMP mice infected for 4 weeks (**Figure 8B, top row**) and 6 weeks (**Figure 8B, bottom row**) are arrayed based on their overall mean intensity. The percentage of pixels within each of the 4 bins is presented as a stacked plot, with the AmyloQuant generated thumbnails representing spatial heatmaps of AG intensity at 5 cyst intervals as presented below each stacked plot. This analysis reveals that despite the mean PAS intensities not being statistically significant (**Figure 8A**), several ΔTgLaf cysts present with varying levels of high intensity (red) PAS labeling. Notable here is that with this cohort of cysts, effective complementation of AG levels is not observed.

As expected for wild type Week 6 tissue cysts (14), markedly increased AG accumulation is evident compared to Week 4 cysts (**Figure 8B**). This phenotype is grossly exaggerated in the case of ΔTgLaf tissue cysts (**Figure 8B**), displaying the expected starch excess phenotype. Even so, not all ΔTgLaf cysts present uniformly high levels AG based on PAS intensity (**Figure 8B**). Notable here, the levels and distribution of AG levels within the COMP line appear to be intermediate to that observed for the WT and ΔTgLaf, pointing to the plasticity of the phenotype.

Several recent studies targeting activities connected with AG metabolism report differences in cyst yield as well as cyst size (21–23, 41). We therefore examined if the loss of TgLaf resulted in any significant impact of cyst size. Contrary to other reports however no differences in cyst diameter were noted for either week 4 or week 6 harvested tissue cysts (**Figure S3A**). Furthermore, we found no strong correlation between cyst size and mean AG levels as defined by PAS labeling intensity (**Figure S3B**).

### Aberrant AG accumulation in **Δ**TgLaf tissue cysts promotes bradyzoite death

The inherent harshness of PAS labeling causes differential labeling intensity related issues with regard to the labeling of bradyzoite nuclei (14) as well as labeling for TgIMC3 a marker of recent replication (14). As a result, the relationship between AG levels and replication associated outputs (packing density and recency of replication) could not be reliably implemented in the context of the TgLaf mutant. We therefore resorted to the direct examination of purified tissue cysts by transmission electron microscopy.

We adapted a protocol designed to capture and image low abundance cells by TEM by making it compatible with our tissue cyst purification protocol (see Methods) (71, 72). TEM imaging revealed that while WT parasites formed largely normal/canonical AGs *in vivo* as seen *in vitro* (**Figure 9** [compare with **Figure 5A**]), ΔTgLaf parasites contained almost exclusively aberrant AGs that mirrored the same morphological defects seen *in vitro* (**Figure 9** and **Figure S4**). ΔTgLaf AGs were irregularly sharpened with a flat, multi-lobed appearance. Importantly, COMP parasites neither over-accumulated nor formed aberrant AGs, demonstrating that this defect is specific to loss of TgLaforin (**Figure 9**).

**Figure 9.**
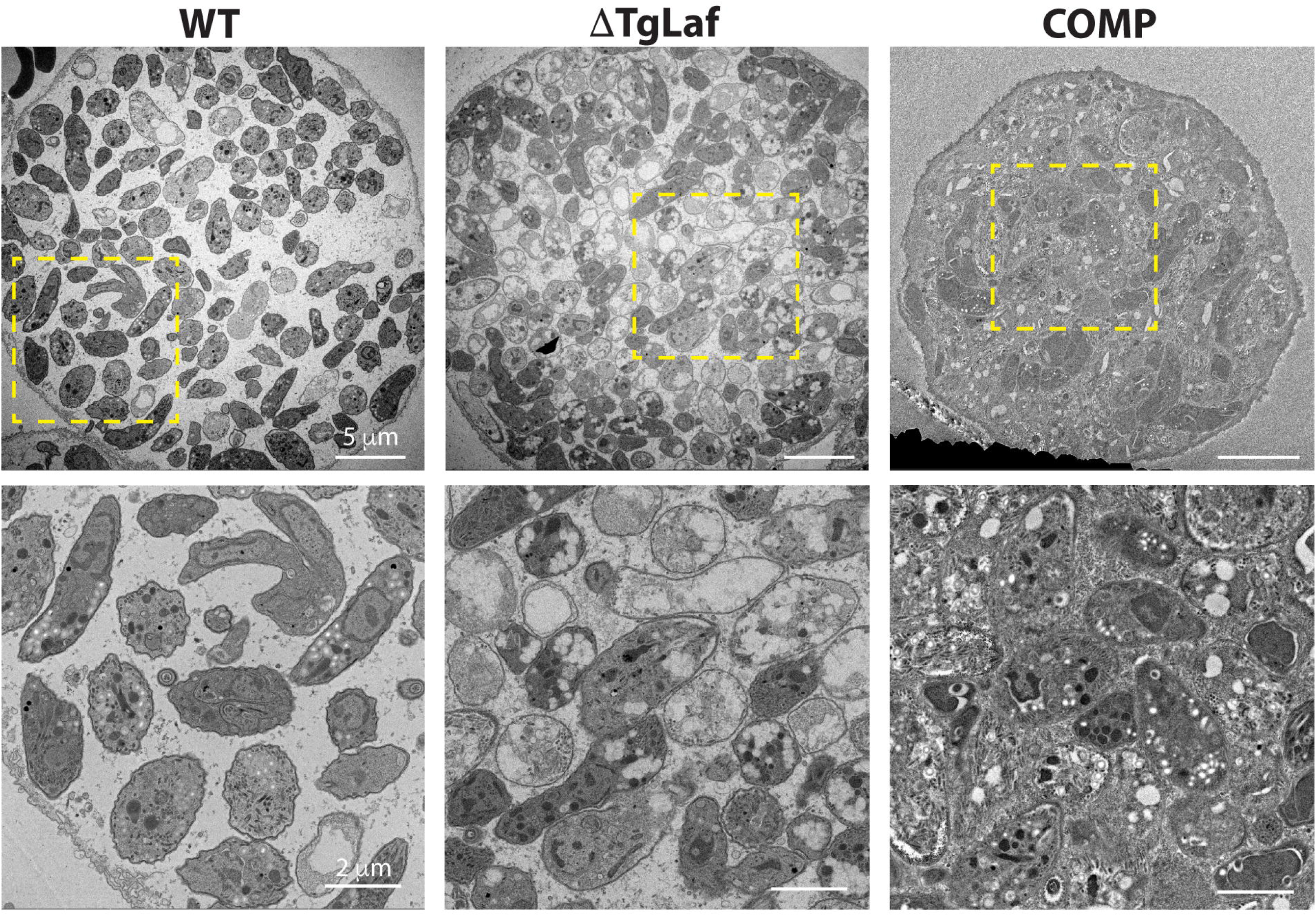
Transmission electron microscopy of purified tissue cyst from infected mouse brains confirms the presence of aberrant AG accumulation within (TgLaf cysts. The accumulation of amylopectin granules within the cytoplasm of WT bradyzoites with parasites exhibiting different levels. Additionally, evidence of active endodyogeny is present. (TgLaf tissue cysts contain a mix of bradyzoites with expected cytoplasmic and organellar contrast as well as others with grossly exaggerated AG levels that and the apparent loss of both the cytoplasmic and organellar contents. The COMP line exhibits AG accumulation levels similar to that observed in WT parasites with the tissue cyst itself appearing to be very tightly packed, with high levels of granular material between individual bradyzoites. Upper panels: scale bar = 5 μm; lower panels (zoom of boxed region from upper panel): scale bar = 2 μm.

In addition to containing aberrant AGs, the internal morphology of ΔTgLaf parasites appeared to be altered by the presence of the AGs, as significant organelle displacement was also noted (**Figure 9, S4**). Many of the ΔTgLaf parasites also appeared hollow or “ghost-like” in appearance resulting from their unstained/absent cytoplasm, which included the apparent loss of the nucleus in several bradyzoites (**Figure 9, S4**). These data suggest that a significant number of the ΔTgLaf bradyzoites were inviable within the cyst (**Figure 9, S4**). Together, these direct and selective impacts of TgLaforin’s loss on both bradyzoite viability and growth *in vivo* establish TgLaforin as a potentially druggable target.

## DISCUSSION

The asexual life cycle of *Toxoplasma gondii* is defined by two fundamentally distinct forms: the rapidly replicating tachyzoite and the slowly growing bradyzoite. These forms represent distinct physiological states that can be further subdivided, particularly within encysted bradyzoites (73). Insights into these physiological states, inferred from transcriptomic analyses, are reinforced in emerging metabolomic studies (74, 75). In these studies, glucose and glutamine, which are both linked to energetics, biosynthesis, and intermediary metabolism, appear as critical metabolites. Importantly, glucose can be stored in polymers like glycogen and amylopectin during times of low energy need. Stored glucose can be present in an accessible and labile form for rapid mobilization such as glycogen, or in a less soluble forms as AGs from which it can be accessed more slowly (76, 77). Reversible glucan phosphorylation facilitates the breakdown of such glucan polymers by disrupting the crystalline helices on the glucan surface (78). *T. gondii* encodes the capacity for reversible glucan phosphorylation (27, 36, 45, 46). The contribution of this process to tachyzoite and bradyzoite biology was evaluated through targeted manipulation of the glucan phosphatase, TgLaforin (36).

The accumulation of AGs within bradyzoites and their apparent absence in tachyzoites has been used as discriminator between these life cycle stages (16, 79). Detailed examination, however, presents a considerably more nuanced picture, alongside emerging evidence that points to rapid glucan turnover within tachyzoites (20). In the current study, using an IgM monoclonal antibody (IV58B6) that specifically recognizes glycogen-like glucose polymers (47, 48), we demonstrate that the stored glucan within tachyzoites is structurally closer to animal glycogen (47, 48) than the insoluble plant-like AG granules found in bradyzoites (17, 19). This duality between life stages may be additionally reflected in the observation that TgLaforin, the glucan phosphatase, and TgGWD (TGME49_214260), the partner kinase that is predicted to phosphorylate *T. gondii* glucan polymers, trace their structural lineages to animals and plants, respectively (45, 46). The glycogen-like glucan polymer appears to be specific to tachyzoites as its levels decrease upon *in vitro* differentiation while the overall PAS intensity increases (**Figure 1A**). This suggests that the tachyzoite glucan and bradyzoite AG are architecturally distinct polymers with respect to both branching frequency, solubility, and phosphorylation status (**Figure 2D**). The glucose-based nature of both particles is supported by the elimination of both IV58B6 and PAS staining with α- amyloglucosidase treatment (**Figure 1B**).

To address the contribution of stored glucans in both tachyzoites and bradyzoites, we disrupted the glucan phosphatase TgLaforin. This enzyme preferentially removes phosphate groups from the C3 carbon on glucose facilitating access to enzymes that release glucose (36). The loss of other glucan phosphatases such as SEX4 in *A. thaliana* and laforin in mammals is accompanied by excessive accumulation of aberrant starch and hyperphosphorylated glycogen in plants and animals (37, 39, 40, 80, 81). Surprisingly, given these penalties in other systems, ΔTgLaf tachyzoites exhibited no gross morphological changes in glucan content, consistent with a recent study in which TgGWD was knocked out (27). These observations contrasts with other KO studies of glucan pathway associated proteins such as CDPK2 (20), glycogen phosphorylase (41), the PP2A holoenzyme (44), and α-amylase (22)which all reported exaggerated glucan accumulation in tachyzoites.

Despite the absence of glucan accumulation in ΔTgLaf tachyzoites, the loss of this gene exerts an effect on tachyzoite glucan metabolism. Consistent with the metabolic defects associated with the loss of laforin in humans (60), loss of TgLaforin in *T. gondii* resulted in altered central carbon metabolism that manifested as ΔTgLaf parasites’ dependence on glutamine (**Figure 4, S2D,E**). ΔTgLaf tachyzoite dependence on glutamine supports recent studies demonstrating that tachyzoites utilize storage glucans for glucose allocation (23) because the presumed loss of efficient glucan degradation results in tachyzoite dependence on glutamine (**Figures 4B,C**). As *T. gondii* tachyzoites primarily utilize glucose and glutamine to support their rapid growth, this dependence on glutamine reinforces previous observations that glutamine can substitute for glucose in this altered metabolic landscape (62, 82). These results are consistent with many previous studies that disrupt glucose and/or glucan metabolism, but contrast with others. Disruption of TgGT1 (TGME49_214320), the only plasma-membrane glucose transporter in *T. gondii* (62, 83), or TgHK (TGME49_265450), the *T. gondii* hexokinase, resulted in upregulation of gluconeogenesis, and parasite growth was highly attenuated with glutamine depletion (82). Most strikingly, parasites lacking starch synthase (TgSS; TGME49_222800) displayed no dependence on glutamine and, in fact, grew faster than WT parasites when both glucose and glutamine were removed from the culture media (23). Interestingly, however loss of TgSS did result in lower glucose flux through glycolysis (23), consistent with our findings that demonstrate a role for the tachyzoite glucan in glucose allocation. This finding may suggest that while the absence of AG in the ΔTgSS parasites may not be detrimental, overaccumulation as observed in ΔTgLaf cysts can contribute to toxicity (**Figure 9**). Perturbations of other glycolytic enzymes also demonstrated varied effects related to the presence of glutamine: loss of the glycolytic enzyme glyceraldehyde-3-phospahte dehydrogenase (GAPDH1) could be rescued with high levels of glutamine (84) (84), but glutamine could not rescue pyruvate kinase (TgPYK1) knockdown parasites (85). Our data indicate that loss of access to key nutrients such as glucose and glutamine has a profound impact on the repeated rounds of the lytic cycle without being attributable to one specific process within the cycle, suggesting that the penetrance of the phenotypic defect manifests cumulatively over time, rather than being hard wired in each infection cycle (**Figures 4D,E, S2D,E**).

Despite glucan metabolism being historically viewed as being important in the chronic infection, TgLaforin protein expression decreased during the tachyzoite to bradyzoite conversion *in vitro* even though its transcript levels do not change (**Figures 2C,D**). This could be a transient observation as the downregulation of glucan catabolism during conversion would facilitate accumulation of AGs for the chronic infection. We therefore examined how the loss of TgLaforin affected the capacity of ΔTgLaf parasites to differentiate *in vitro*. The ΔTgLaf parasites exhibited no defect in AG-accumulation kinetics, detected by PAS staining, or in cyst wall formation, detected with DBA lectin over the course of the in vitro conversion assay (**Figure 5A, B**). The lack of difference in PAS labeling between both the WT and the ΔTgLaf lines, however, did not reveal the differences noted by TEM (**Figure 5C-E**). As initially hypothesized would be the case in both tachyzoites and bradyzoites, loss of TgLaforin resulted in aberrant AG accumulation within *in vitro* bradyzoites that is marked by changes in both level and morphology (**Figures 5C-E**), as seen in plants and vertebrates (37–40, 81). AGs in the ΔTgLaf parasites were not only present at higher levels but were potentially chemically distinct considering their differential binding to TEM contrast metals (**Figure 5E**). Given that TgLaforin is a confirmed glucan phosphatase (36), we speculate that AG hyperphosphorylation may account for both altered morphology and appearance by TEM.

These context-specific phenotypes suggested that the ΔTgLaf mutant would manifest phenotypic differences in both the acute and chronic phases on infection *in vivo*. Indeed, the loss of TgLaforin was associated with a markedly reduced symptomology and associated mortality compared to both the parental and complemented parasites during acute tachyzoite-initiated infection (**Figures 6A,B**). Not only was there a delay in symptomatic disease, but also a reduction in disease severity and overall cyst burden. Symptomology in the acute infection is driven by an increasing parasite burden driving an overexuberant host inflammatory response (4, 86). The delayed symptom onset suggests growth inhibition by the stringent *in vivo* environment that more effectively controls ΔTgLaf parasite infection with less robust inflammation. Notably, the delayed and milder course of the tachyzoite infection resulted in a lower overall cyst burden in surviving animals compared to infection with both WT and COMP parasites (**Figure 6C**).

Infection with ΔTgLaf tachyzoites resulted in fewer tissue cysts being generated relative to WT and COMP parasite (**Figure 6C**) The basis for this is not clear, but suggestive of a regulatory imbalance that is masked in tachyzoites but evident in tissue cyst-initiated infections which have the additional burden of converting from bradyzoites to tachyzoites, surviving the chronic infection before forming new tissue cysts. This could potentially parallel the observations with plaque formation in vitro (**Figure 4, S2**).

Our ability to quantify and map relative AG levels within tissue cysts using AmyloQuant (14), confirm that the loss of TgLaf does not fundamentally alter the initiation and early progression of an AG cycle as evidenced by the effect of the mutation not being evident in Week 4 tissue cyst in the form of the expected starch excess phenotype (**Figure 8**). The effect of the ΔTgLaf mutation becomes evident in week 6 tissue cysts where a massive increase in AG accumulation (**Figure 8**) is likely due to an imbalance caused by the predicted defect in AG turnover. As with other phenotypes associated with this mutation, the effect continues to exhibit phenotypic variation noted by roughly a third of the imaged cysts lacking a significant proportion of high intensity pixels (red), while others are completely oversaturated (**Figure 8**). These distinct populations are not reflected in any way based on the size of the cyst (**Figure S3**). In fact, contrary to other reports regarding AG metabolism associated genes, the TgLaf mutation has no significant effect on tissue cyst size (**Figure S3**).

While we were not able to accurately quantify either the nuclear number or TgIMC3 intensity distributions as markers of packing density and replicative activity in PAS stained tissue cysts (14), TEM analysis exposed the true consequence of the loss of TgLaf within tissue cysts in vivo (**Figure 9, S4**). Wild type tissue cysts presented with clusters of small amylopectin granules as well as evidence of active endodyogeny (**Figure 9**). In contrast ΔTgLaf cysts presented bradyzoites laden with AG to the point where other organellar structures were obscured (**Figure 9, S4**). Several bradyzoites lacked nuclei while others lacked the cytoplasm (**Figure 9, S4**). Together, these features are incompatible with viability. The presence of such features does not apply to all bradyzoites within the cyst accounting for the fact that these cysts are able to initiate a new infection (**Figure 7**.), albeit with lower virulence and cystogenic potential. These finding suggest that AG metabolism is under tight control as dysregulated accumulation can result in cumulative defects resulting in toxicity and death. The high frequency of these abnormal parasites suggests that reversible glucan phosphorylation and TgLaforin specifically represent legitimate bradyzoite specific drug targets. We recently described a small molecule that inhibits recombinant TgLaforin (36) which serves as a potential starting point in the development of a new class of anti-*Toxoplasma* therapeutic agents. Particularly exciting in this context is the fact that a class of drugs exhibiting efficacy with tissue cyst clearance (atovaquone (87, 88) endochin-like quinolones (89–91) and JAG21 (92)), all target mitochondrial respiration. When glucose is limiting, mitochondrial respiration can be driven by glutamine. This provides an opportunity for combination therapy to promote the clearance of toxoplasma tissue cysts as a means of mitigating the risk of reactivation.

## METHODS

### Fibroblast and parasite culture and maintenance

All parasite lines were maintained in human foreskin fibroblasts (HFFs; ATCC) in Minimal Essential Media-α (MEM-α; Gibco) supplemented with 7% heat-inactivated fetal bovine serum (FBS; Gemini Bio), 100 U/mL penicillin, 100 μg/mL streptomycin, and an additional 2 mM L-glutamine (Gibco; 4 mM total L-glutamine). Cells and parasites were incubated at 37°C and 5% CO_2_ in a humidified incubator. Genetically modified parasites were maintained in MEM-α containing 7% dialyzed FBS (Gemini Bio) and either pyrimethamine (1 μM), mycophenolic acid/xanthine (MPA: 25 μg/mL, xanthine: 50 μg/mL), or 6-thioxanthine (6-Tx: 80 μg/mL).

Assays analyzing the effects of glutamine deprivation used Dulbecco’s Modified Eagle Medium (DMEM). Both glutamine-replete (Gibco, 11966025) and depleted (Gibco, 11054020) DMEM were supplemented with 7% dialyzed FBS. Glutamine-replete media from the supplier lacked other key nutrients and was modified to contain 5 mM glucose, 1 mM sodium pyruvate, and 4 mM L-glutamine.

### Generation of T. gondii mutant lines

*Type II ME49ΔHXGPRT* (“WT”—the parental line utilized to generate all other lines in this study): This line was generated in a previous study using CRISPR/Cas9 targeting of TgHXGPRT and selection with 6-Thioxanthine (52).

*TgLaforin-3xHA-HXGPRT*: TgLaforin was epitope tagged with HA at the C-terminus using CRISPR-Cas9 to disrupt the TgLaforin 3’UTR immediately downstream of the endogenous stop codon as has been previously described (53). Briefly, a sgRNA immediately downstream of the TgLaforin stop codon was designed using the EuPaGDT design tool (http://grna.ctegd.uga.edu). The top hit was selected (**Table S1**) and used to replace the sgRNA sequence in pSAG1::CAS9-U6::sgUPRT, a plasmid containing both Cas9-green fluorescent protein (GFP) and an interchangeable sgRNA scaffold (56); (**Table S2**). Replacement of the interchangeable sgRNA was accomplished using a Q5 site-directed mutagenesis kit (**Table S3**) (New England BioLabs). The TgLaforin-HA tagging construct was generated by amplifying the 3’ end of the TgLaforin-HA construct generated for complementation (see generation of COMP line below and **Tables S2 and S3**) along with the connected HXGPRT selectable marker. Both the TgLaforin-HA PCR-amplicon and the CRISPR-Cas9-GFP were transfected into 1.4×10^7^ *T. gondii* ME49ΔHXGPRT parasites (2:1 insert:plasmid molar ratio; 30 μg DNA total) by electroporation with a time constant between 0.16 and 0.20 msec (BioRad Gene Pulser II). After 24 h, surviving parasites were syringe-passaged from infected HFFs with a 27 G needle to lyse host cells, and gravity-filtered through a 10 μm filter to remove host-cell debris. Successful transformants were then enriched by use of fluorescence-activated cell sorting (FACS; Sony SY3200, installed in a biosafety level II cabinet) to select parasites expressing Cas9-GFP from the transfected plasmid by isolating GFP+ parasites. HFFs were infected with GFP+ parasites, and then placed in media containing MPA/xanthine 24 h later to select for restoration of HXGPRT. MPA/xanthine-resistant parasites were cloned by limiting dilution into a 96 well plate. Wells containing single plaques were picked 7 days later and expanded. Genomic DNA was extracted from clones using a Proteinase K treatment detailed elsewhere (93). Successful tagging of TgLaforin was verified using sequencing, immunoblotting, and IFA.

*ME49ΔHXΔTgLaforin* (“ΔTgLaf”): TgLaforin was disrupted using a CRISPR-Cas9 mediated strategy as detailed above, with several differences. Briefly, a single sgRNA was designed to target the first exon of TgLaforin with the top hit from EuPaGDT (**Table S1**). To disrupt TgLaforin with a selectable drug marker, DHFR-TS*, a pyrimethamine-resistant mutant of the DHFR gene, containing a 5’-NcGra7 promotor and DHFR 3’UTR was amplified from pJET-NcGra7_DHFR (**Table S2**). Amplification utilized primers containing 40 nt extensions homologous to the 5’- and 3’-UTR of TgLaforin to encourage homologous recombination-mediated whole-gene replacement with the drug cassette (**Table S3**). Both the PCR-amplified DHFR* homology cassette and the CRISPR-Cas-GFP plasmid were transfected and FACS-sorted as described above. GFP+ parasites underwent drug selection in pyrimethamine. Parasites were then cloned and expanded as detailed above. Successful integration of the DHFR* cassette into the TgLaforin locus was verified using PCR with inside/out primer pairs to the chimeric, interrupted gene (**Table S3**). Loss of TgLaforin transcription was verified by purifying RNA from TgLaforin clones on RNeasy spin columns (Qiagen). Using the Promega Reverse Transcriptase System, cDNA was synthesized from RNA extracts. Primers designed for full-length TgLaforin amplification were then used to verify loss of TgLaforin cDNA in knockout lines.

*ME49ΔHXΔTgLaforin+ChrVI-TgLaforin* (“COMP”): Complementation of TgLaforin was also executed using a CRISPR-mediated strategy. A sgRNA to a neutral locus on chromosome VI identified previously (58) was generated using the same mutagenesis strategy as above (**Table S1 and S3**). A full length TgLaforin cDNA containing its endogenous 5’UTR (2000 bp upstream from gDNA) was synthesized by GenScript and inserted into a pHA3x-LIC vector (**Table S2**) containing a C-terminal HA tag and a DHFR 3’UTR, linked to the HXGPRT selectable marker (named “TgLaforin-HA3x-LIC”; also used above for endogenous tagging to create the TgLaforin-HA line). The entire construct (5’UTR:TgLaforin-cDNA:DHFR-3’UTR:HXGPRT) was amplified from the vector and co-transfected into ΔTgLaf parasites with the CRISPR-Cas9 plasmid as done above. Successful transformants that received the HXGPRT marker were selected with MPA/xanthine. Successful insertion of TgLaforin along with its promoter was verified using PCR (**Table S3**), immunoblotting, and IFA with an anti-HA antibody (Abcam).

### Immunofluorescence (IF) staining

HFFs were grown on glass coverslips until confluent and subsequently infected. Infected HFFs were fixed with either methanol (MeOH) (100%, -20°C) or methanol-free paraformaldehyde (PFA) (4% in phosphate-buffered saline (PBS); Electron Microscopy Sciences) as indicated below for each antibody. Infected HFFs fixed with PFA were permeabilized in 0.1% TritonX-100 in PBS++ (PBS containing 0.5 mM CaCl_2_ and 0.5 mM MgCl_2_) for 10 min at room temperature (RT). Primary and secondary antibodies were diluted in 3% (w/v) bovine serum albumin (BSA; Fisher) in PBS++. Samples were first incubated with the primary antibody (αHA-1:1,000; αSAG-1:10,000; αGAP45- 1:5,000; αGRA3-1:1500; IV58B6-1:50) at RT for 45 min, washed 3x with PBS++, and then incubated with fluorescent secondary antibodies (1:2,000) and 4’,6-diamidino-2- phenylindole (DAPI; 300 nM) for 45 min. Secondary antibodies (Invitrogen) were conjugated to either Oregon Green or Texas Red fluorophores and specific to the species and class of primary antibody used. Samples were then washed 3x with PBS++ before mounting the coverslip on a glass slide using MOWIOL mounting media.

Immunofluorescence staining was visualized using a Zeiss AxioVision upright microscope with a 100X 1.4 numerical-aperture oil immersion objective, and images were acquired using a grayscale Zeiss AxioCam MRM digital camera. Grayscale images were pseudo-colored in ImageJ using magenta (Texas Red), yellow (Oregon Green), and cyan (DAPI), and further alterations to brightness and contrast were also made in ImageJ when deemed appropriate. For all assays in which staining intensity was compared across treatments and parasite lines, concentrations of antibodies, exposure times, and alterations to brightness/contrast were identical. Colocalization of fluorescent antibodies/reagents was quantified using Pearson’s coefficient calculated with the JACoP plugin on ImageJ (94).

### *PAS staining*-tachyzoite and in vitro bradyzoite

PAS staining of tachyzoites and in vitro bradyzoites was done on infected HFFs fixed in 4% PFA and permeabilized as above. Coverslips were then washed 3x in tap water before the addition of 1% periodic acid (Sigma-Aldrich) for 5 min. Coverslips were then washed with three changes of tap water. Schiff’s reagent (diluted 1:4 in tap water) was added for 15 min. Coverslips were subsequently washed 10x with tap water to develop stain before being incubated with DAPI for 10 min and then mounted as above. PAS-stained samples were visualized using fluorescence microscopy (excitation: 545 nm, emission: 605 nm). When PAS was co-stained with antibodies, primary antibodies were incubated with PAS-stained slides overnight in BSA at 4 °C before standard secondary staining.

PAS labeling of methanol fixed tissue cysts was performed as described elsewhere, using Schiff reagent diluted 1:10 in tap water as described elsewhere ().

Samples treated with acid-α-amyloglucosidase (GAA) (from *Aspergillus niger*, >260 U/mL, Sigma) were incubated with GAA after permeabilization. GAA was diluted 1:50 in 50 mM sodium phthalate buffer, pH 5.5, and samples were treated for 24 h at room temperature. Untreated controls were incubated in phthalate buffer without GAA. Samples were then stained with PAS or IV58B6 as described in the IF-staining workflow above.

### In vitro bradyzoite conversion assay

Tachyzoites were converted to bradyzoites *in vitro* using alkaline stress as has been done previously with several modifications (95). HFFs grown were infected with tachyzoites in standard cell culture media. 4 h later, media was replaced with RPMI 1640 (Gibco 31800022) supplemented with 50 mM HEPES and adjusted to pH 8.2 with NaOH. Parasites were then cultured for 2-6 days at 37°C, ambient CO_2_, and sealed in Parafilm. Media was replaced every other day to maintain the basic pH. Parasites were fixed in PFA and stained with fluorescein conjugated *Dolichos biflorus* agglutinin (DBA; 1:1000, Vector Laboratories) and PAS. Images were obtained in grayscale on a Zeiss AxioVision upright microscope as described above. To determine the degree of labeling with DBA or PAS, the Fiji/ImageJ (96) was used to create a binary mask outlining cysts that was applied to the PAS-stained image to measure the greyscale intensity of each ROI (i.e. each individual vacuole/*in vitro* cyst).

### Transmission electron microscopy of in vitro tachyzoites and bradyzoites

Transmission-electron microscopy (TEM) was performed as done previously (97). Blocks were stained at the University of Kentucky’s Imaging Center in the College of Arts and Sciences. Blocks were trimmed and sectioned on an ultramicrotome with a diamond knife. Sections were placed on copper grids and then contrast stained with lead citrate. Micrographs were collected at the University of Kentucky’s Electron Microscopy Center on a Talos F200X TEM (Thermo) operated at 200 kV accelerating voltage with a 50 μm objective aperture inserted to enhance contrast using a 16M pixel 4k x 4k CMOS camera (Ceta, Thermo Scientific). AG size and grayscale values were measured in ImageJ.

### Immunoblotting

Parasites were syringe lysed from host cells, pelleted, and 2×10^6^ parasites were resuspended in SDS-PAGE sample buffer and boiled for 10 min before being run on a single lane of a 10% polyacrylamide gel. The gel was then transferred to a 0.2 μm PVDF membrane (BioRad) using a Turbotransfer System (BioRad) for 7 min at 25 V. The PVDF membrane was blocked in 5% (w/v) non-fat milk in Tris-buffered saline plus Tween-20 detergent (TBST; 0.1% Tween-20) for 20 min before being probed with a primary antibody (αHA-1:1,000; αGAP45-1:5,000; αSRS9-1:1,000; αSAG1-1:10,000) in non-fat milk overnight at 4°C (Cell Signaling C29F4). The blot was washed 3x with TBST before probing with either HRP-conjugated α-rabbit or α-mouse-IgG (Jackson Laboratories). Blot was washed and developed for 5 min using SuperSignal^TM^ West Pico PLUS (Thermo Scientific) and visualized on a GelDoc station (BioRad).

### Steady state polar metabolite analysis

Parasites were prepared as previously described (61). Confluent HFFs were infected with parasites at a multiplicity of infection (MOI) of 2 to achieve a high density of parasites after 48 h of growth (>80% cells containing >32 parasites each). Plates containing infected HFFs were placed on ice, media removed, and the monolayer was washed 2X with ice-cold PBS. Parasites were harvested on ice in a 4 °C cold-room. Cells were scraped from plate surface, resuspended in PBS (8 plates/50 mL PBS), and centrifuged at 1000*g* for 10 min at 4 °C. PBS was removed, the cell pellet was resuspended in 2 mL PBS, and syringe passaged successively in 23 G and 27 G needles. The soluble host cell lysate was removed by centrifugation (1000*g*). The pellet was resuspended in 5 mL PBS and host-cell debris was removed by syringe-filtering the suspension through a 3 μm filter (Whatman). Filtered parasites were then pelleted, resuspended in 1 mL PBS, and counted on a hemacytometer. Parasites were pelleted a final time at 14,000*g* for 30 s at 4 °C, supernatant was removed, and pelleted parasites were flash frozen in liquid nitrogen and stored at -80 °C until metabolite extraction.

#### Polar metabolite extraction

Polar metabolites were extracted in 0.5 mL -20 °C 50% methanol (MeOH) containing 20 μM L-norvaline (procedural control) for 30 min on ice. During the 30 min incubation, samples were regularly vortexed. Samples were then centrifuged at 14000*g* for 10 min to pellet insoluble material (protein, DNA, RNA, and glycans). Supernatant containing polar metabolites and pellet were dried separately on a SpeedVac (Thermo) at 10^-3^ mBar until methanol (MeOH) was completely sublimated and only dried pellet remained.

#### Pellet hydrolysis and extraction

Dried fraction containing protein was hydrolyzed by resuspending the pellet in 2 N HCl (final concentration) at 95°C for 2 h. Hydrolysis was quenched, and hydrolyzed amino acids were extracted by the addition of an equal volume of 100% MeOH with 40 μM L-norvaline such that the final concentration was 50% and 20 μM, respectively. Extraction and drying then proceeded as described above.

#### Sample derivatization

Dried samples (both polar metabolites and hydrolyzed protein) were derivatized in 70 μL 20 mg/mL methoxyamine hydrochloride in pyridine for 90 min at 30 °C. Samples were then centrifuged at 14000*g* for 10 min to remove any particulate, and 50 μL of the methoxyamine supernatant was mixed with 80 μL *N*- methyl-*N*-trimethylsilyl trifluoroacetamide (MSTFA) and incubated for 30 min at 37°C. Samples were then transferred to amber glass chromatography vials and analyzed by GC/MS.

#### GC/MS analysis

Metabolites were analyzed on an Agilent 7800B GC coupled to a 5977B MS detector using a previously established protocol (98). Automated Mass Spectral Deconvolution and Identification System (AMDIS) was used to analyze metabolites by matching metabolites to the FiehnLib metabolomics library via retention time and fragmentation pattern. Quantification of metabolite levels was performed in Mnova. Sample abundance was normalized to L-norvaline (procedural control) and protein from the protein pellet (experimental control). Steady state metabolites are presented as the mean of three independent replicates.

### Conventional plaque assays

HFFs were grown in 12-well plates until confluent. HFFs were subsequently infected with 200 parasites/well under standard cell culture conditions. Wells were washed with PBS to remove residual invasion media, and media was changed to glutamine replete or depleted media 4 h post-infection to allow for invasion. Plates remained undisturbed for 10 days before the infected HFFs were fixed with 100%, -20°C MeOH for 20 min, stained with 1% crystal violet solution for 20 min, and then the plaques were de-stained with repeated tap water washes. Zones of lysis (white clearings) could be visualized against intact cells (purple). Images of plaques were obtained by scanning plates on an Epson Perfection V600 photo scanner at a resolution of 600 dpi. The plaques were measured by pixel area using ImageJ. Plaque assays were conducted on three independent replicates, and plaque size from these experiments were aggregated to highlight variability in plaque sizes.

### Modified plaque assays

A modified plaque assay was developed to visualize foci of infection at higher magnification where zones of clearing were not readily evident. Confluent HFF monolayers on glass coverslips in 24 well plates were infected with WT, ΔTgLaf, and COMP parasites in replete media for 4 hours to allow for invasion. The monolayers were washed gently 3 times with PBS and either fresh replete media or glutamine-depleted media were added for 3 or 6 days to appropriate wells. Infected monolayers were washed and fixed with MeOH (-20 °C) and subjected to IF using DAPI and GRA3, an antibody that detects the PVM. Individual plaques were imaged on coded blinded slides using a 10X objective, and their perimeters and encompassed areas were measured using Image J. Host cell clearance within plaques was similarly measured, and the extent of clearance was represented as a percentage for each individual plaque. A total of three independent replicates were performed.

### Egress assays

HFFs were grown to confluency in 35 mm glass bottom dishes (MatTek, P35G-0-14-C). Two days before infecting HFFs on glass bottom dishes, both HFFs and parasites were independently pre-treated in either glutamine-replete or -depleted media (see above for media formulations). After 48 h pre-treatment, 10^5^ parasites of each line (WT, ΔTgLaf, and COMP) in each pre-treatment (gln+/-) were added to fresh dishes and allowed to grow for 48 h so that most vacuoles contained >32 parasites each. Several hours before egress, media in each infected plate was adjusted to 1.5 mL and allowed to equilibrate at 37°C in 5% CO_2_. The calcium ionophore A23187 (Cayman Chemical Company) was prepared as a 2 mM stock in DMSO and diluted in (+/-) gln media to make a 4X concentration of 12 μM and maintained at 37°C throughout the assay. Zaprinast was likewise prepared as a 100 mM stock in DMSO and diluted into media at a 4X concentration of 2 mM. Egress was triggered by the addition of 0.5 mL 4X A23187 to infected HFFs (3.0 μM final concentration) or 0.5 mL 4X-zaprinast (500 μM final concentration). Egress was monitored on a Nikon Eclipse Ti2 inverted microscope with a 40X phase air objective modified with a 1.5X optivar. Several fields containing vacuoles were selected from each plate, and an image was obtained 10 s after triggering egress from each field once every 5 s for 5 mins (61 images/field) on a Nikon DS-Ri2 color camera. Videos of each field were assembled on NIS Elements software. Egress was monitored using standard deviation of pixel intensity and determined by inflection point of change in standard deviation of pixel intensity. Inflection point was calculated by fitting a gaussian curve to the first derivative of the standard deviation in pixel intensity and calculating the mean of the curve. Technical replicates (fields on each plate) were averaged for each biological replicate (average of fields from each plate).

### Replication assays

HFFs were grown on glass coverslips in a 24-well plate until confluent. Two days before infecting HFFs on coverslips, both HFFs and parasites were independently pre-treated in either glutamine-replete or -depleted media. After pre-treatment, 10^4^ parasites of each line (WT, ΔTgLaf, and COMP) in each condition (gln+/-) were added to 3 coverslips each. 24 hours later, infected HFFs were fixed in MeOH and stained with Rb-α-SAG1 (1:10,000) and DAPI for ease of visualization. Counting of parasites/vacuole was performed for each line/condition on coded blinded slides with the identity of samples revealed upon completion of the counting. A total of three independent replicates were performed.

### Mouse infection studies

4- to 6-week old CBA/J mice of both sexes (Jackson Laboratories, Bar Harbor, ME) were injected intraperitoneally (i.p.) with either 100 WT, ΔTgLaf, or COMP tachyzoites, or with 20 tissue-cysts from brain homogenates derived from previously infected mice. In either case, parasites/cysts were suspended in a final volume of 0.2 mL serum-free, Opti-MEM media (Gibco). Mice were then monitored and assigned a body index score daily. Monitoring frequency increased to twice a day once symptomatic throughout the course of infection as previously described (52). When symptomatic, mice were administered a gel diet and wet chow on the cage floor and given 0.25-0.5 mL saline solution subcutaneously as needed. Moribund mice were humanely euthanized. Euthanasia of both moribund mice and mice sacrificed at the time of tissue cyst harvest was performed by CO_2_ asphyxiation, followed by cervical dislocation. The number of mice used for each experiment is indicated within relevant figures. All protocols were carried out under the approval of the University of Kentucky’s Institutional Animal Care and Use Committee (IACUC).

### Tissue cyst purification

Tissue cysts were purified as previously described using discontinuous Percoll gradients (11, 70). Processing of two sex/infection-matched brains was performed on each gradient. Cysts were collected in 1 mL fractions from the bottom of the centrifuged Percoll gradient using a peristaltic pump adjusted to a flow rate of 2 mL/min. To quantify tissue cysts, 10-20 μL of each fraction was placed into 100 μL PBS in the well of a 96- well plate, pelleted and directly enumerated at 20X magnification in each well. Total cysts per mouse were calculated by summing the total number of cysts in each fraction and dividing the total by two to adjust for brain homogenization in pairs. Each pair of mice was presented as a single averaged data point. Tissue cysts were pelleted onto slides using a Cytospin centrifuge and fixed and stored in 100% MeOH (-20°C) until staining.

### Quantification of AG levels in tissue cysts based on PAS labeling intensity

Tissue cysts deposited on glass slides were fixed and stored in methanol at -20°C. Slides were equilibrated to room temperature and stained with PAS using conditions optimized for staining tissue cysts (14). Cysts were additionally stained with DBA lectin to demarcate the cyst boundary. Thirty randomly acquired cysts were imaged using a fixed exposure as a z-stack with a 0.24μm step, with the center slice used for quantification without deconvolution using AmyloQuant, a purpose developed quantitative imaging based application (14). The distribution of pixel intensities was defined in 4 bins representing the background (black: 0-10 grayscale), low (blue: 10-25), intermediate (green: 25-50) and high (red: >50) proportion of pixels (14). The relative distribution of PAS intensities across the 4 bins is represented using a stacked plot with the distribution patterns arrayed from lowest to highest intensity for each cohort. AmyloQuant additionally presents a spatial heat map to allow for the distribution of PAS intensity (AG) to be revealed within the imaged cyst. The mean pixel intensity of each cyst was additionally determined following the definition of the ROI defined by the boundary of the cyst wall using Image J. Measurements of cyst diameters in microns was achieved using the Zeiss Zen imaging software.

### Preparation of in vivo tissue cysts for TEM imaging

To prepare *T. gondii* tissue cysts generated *in vivo* for TEM, cysts were isolated from the brains of infected mice as detailed above through the counting step. The Percoll fraction containing mouse red blood cells (RBCs) was also recovered. After combining Percoll fractions containing tissue cysts and diluting with PBS to a volume of 15 mL (maximum of two 1-mL fractions were combined before dilution), cysts were pelleted for 15 min at 1000*g* at 4 °C. To maximize cyst recovery, 10 mL supernatant was removed, and the remaining 5 mL was divided into 1 mL fractions for the top 4 mL, and the bottom 1 mL directly above the pellet was sub-fractionated into 100 μL volumes. Typically, the majority of cysts were localized to within 300 μL of the Percoll pellet, rather than in the pellet itself. Sub-fractions containing cysts were once again combined and diluted (typically 200-300 μL diluted with 1-mL PBS) in a 1.5 mL Eppendorf tube, and then pelleted in a swinging-bucket rotor for 10 min at 1000*g* and 4 °C. Leaving the pellet undisturbed, all but 50 μL of the supernatant was removed. A small volume (∼5-10 μL) of the reserved RBC fraction was added to the remaining volume for ease of visualizing the pellet throughout the remaining processing steps.

To ensure the detection of the relatively rare cyst population, a previously described protocol (71, 72) was adapted to concentrate the cysts into a small agarose block. A 1.33X fixative solution of glutaraldhehyde (GA) in cacodylate buffer was prepared containing 4% GA and 133 mM sodium cacodylate. 150 μL fixative solution was then added to the 50 μL sample, bringing the total volume to 200 μL such that the final concentration of GA was 3% and sodium cacodylate was 100 mM. Cysts were then incubated at room temperature for 1 h in fixative. While cysts were in fixative, 4% low-melt agarose (BioRad) was prepared in 100 mM sodium cacodylate buffer and kept liquid at 70 °C until needed. After fixation, cysts were pelleted again at 1000*g* for 10 min at room temperature in a table-top centrifuge. All but 50 μL supernatant was once again removed, and 200 μL warm low-melt agarose was slowly added on top of fixed, pelleted cysts (3.2% agarose, final concentration). Suspension was then centrifuged again at 1000*g* for 10 min at 30 °C to keep the agarose semi-liquid, and then placed on ice for 20 min to solidify agarose. After solidification, entire agarose plug was removed from tube with a small a wooden dowel that had been whittled into a thin scoop. This agarose plug was placed in a Petri dish, and the pellet was carefully cut out of the plug with a razor blade to create a 1 mm^3^ block. The agarose block was then stored in 1X GA/cacodylate buffer overnight at 4°C. Processing of the block from post-stain onward was then identical to TEM processing described above. During sectioning, thick sections cut on a glass knife were stained with toluidine blue and examined for cysts using a light microscope prior to ultra-thin sectioning once the cyst containing later was identified.

## Data analyses

All data analyses, including graph preparation and statistics, were performed using GraphPad Prism 9 or 10. Details on statistical tests applied are presented in the specific figure legends.

## Supplemental Files

**Supplemental File 1.**

**Figure S1.** Schematic of TgLaforin complementation strategy and confirmation of successful expression of TgLaforin. **A.** Schematic of TgLaforin complementation into ΔTgLaf parasites in which a PAM site was chosen at a neutral locus previously identified in chromosome VI (99) to insert TgLaforin cDNA under its endogenous promoter. The TgLaforin construct was connected to the HXGPRT selectable drug marker and inserted using NHEJ. **B.** PCR confirmation of integration of TgLaforin construct into chromosome VI. Primer sets are indicated above amplicons. WT primers amplify the same locus as in Figure 3 (“PCR 1”), also present in the COMP line. KO primers amplify the chimeric locus depicted in Figure 3 (“PCR 2”). Presence of the KO amplicon confirms that KO locus remains intact in COMP line. COMP primers amplify the chimeric locus generated upon insertion of the complementation construct. VI primers amplify the native chromosome VI locus, which is lost only in the COMP line. **C,** Western blot confirms expression of TgLaforin-HA in complemented parasites. Tagged LAF-HA parasites serve as a comparison to confirm the correct MW (62 kDa) and expression level. **D,** IFA demonstrates restoration of cytoplasmic, punctate localization of TgLaforin. Scale bar = 5 μm.

**Supplemental File 2**

**Figure S2.** Loss of TgLaforin results in cumulative defects that cannot be pinpointed to a single aspect of lytic cycle. **A.** Calcium ionophore-stimulated egress assay in which parasites were pre-starved of glutamine for 48 hours, seeded onto HFFs and allowed to grow for 48 hours to produce vacuoles containing >16 parasites, and stimulated with 3 μM A23187. Egress was monitored by video microscopy, and time to egress was monitored as described in Materials and Methods. Data is the average of 3 biological replicates that each consist of 4-5 technical replicates. COMP experiments measured 2 biological replicates. **B.** Zaprinast stimulated egress assay performed as described for ionophore, however 500-μM zaprinast was used to stimulate egress. Data is the average of 3 biological replicates that each consist of 2-5 technical replicates. COMP experiments measured 2 biological replicates. **C.** Replication assay in which parasites were pre-starved of glutamine for 48 hours, re-seeded into HFFs, and counted after 24 hours of growth. Numbers (2, 4, or 8) indicate the number of tachyzoites counted per vacuole. Data is the average of 3 biological replicates with at least 70 vacuoles counted per replicate. **D.** Representative images of plaque formation at days 3 and 6, +/- glutamine. Images were taken at 10X magnification using a SAG1 antibody to visualize vacuoles and developing plaque size. The boundaries of vacuoles/ plaques were traced manually to define their area. Regions of cell clearance was additionally scored. **E,** Percent of plaques cleared was measured at both days 3 and 6 by dividing the area of the clearing by total plaque size. Statistical comparisons were done using an ordinary one-way ANOVA using Tukey’s post-hoc test to correct for multiple comparisons. Error bars depict SD from the mean. Statistical significance is indicated as follows: ns=p>0.05.

**Supplemental File 3**

**Figure S3.** The loss of TgLaforin does not have any impact on distribution of tissue cyst sizes. **A**. Tissue cyst diameters of 30 cysts each per line (WT, ΔTgLaf, COMP) per time point (Week 4 and Week 6) were measured using DBA labeled cyst wall as the delimiter using the Zeiss Zen software functionality. While all lines exhibit considerable variability, no statistically significant differences in cyst size were noted in the DTgLaf mutant relative to WT and COMP cysts. **B**. The relationship between tissue cyst size and AG accumulation defined by mean PAS intensity was not found to have any significant correlation.

**Supplemental File 4**

**Figure S4.** Additional images of *in* vivo tissue cysts. **A.** A WT tissue cyst with two zoomed in areas (orange and yellow dashed line boxes) exhibit variability in the levels of AG and present with well defined cytoplasmic and organellar contents. **B.** A ΔTgLaf tissue cyst presents with a mix of bradyzoites with both cytoplasmic and organelles as well as bradyzoites lacking clearly defined organelles and/or cytoplasm. In addition enucleated bradyzoites are evident (asterisks) as well as the displacement of rhoptries by elevated levels of AG in the cytoplasm. Left column scale bar = 5 μm; right column zoom scale bars = 2 μm.

**Supplemental File 5**

**Supplemental Table 1**. sgRNA sequences used in this study.

**Supplemental File 6**

**Supplemental Table 2.** Plasmid constructs used in this study.

**Supplemental File 7**

**Supplemental Table 3**. DNA primer sequences and their application in this study

## Acknowledgements

The authors wish to acknowledge Dr. Elizabeth Watts who conducted early exploratory studies that encouraged the pursuit of this project, Jim Begley (Imaging Center, University of Kentucky) for preparation of EM samples and willingness to explore new methodologies for isolating rare *in vivo* cysts, Jillian Cramer (Electron Microscopy Center, University of Kentucky) for acquisition of TEM images, and Jennifer Strange (Flow Cytometry and Immune Monitoring Core Facility, University of Kentucky) for assistance with isolation of GFP+ *T. gondii* mutants using flow cytometry.

We additionally thank Dr’s David Sibley (Washington University) and Peter Bradley (UCLA) for their gifts of DNA constructs and antibodies.

## Author contributions

Conceptualization (R.D.M., A.P.S., M.S.G.), Methodology (R.D.M., A.D., C.O.B., A.T., A.P.S., M.S.G.), Formal analysis (R.D.M., C.A.T., J.S.M., L.E.A.Y., R.C.S., A.T, A.P., C.W.V.K., A.P.S.), Investigation (R.D.M., C.A.T., J.S.M., L.E.A.Y., A.P.S.), Resources (M.S.G., A.P.S.), Writing—original draft (R.D.M., A.P.S.), Writing—review and editing (all authors), Visualization (R.D.M., C.A.T., J.S.M., A.P.S.), Supervision (M.S.G., A.P.S.), Project administration (R.D.M., M.S.G., A.P.S.), Funding acquisition (R.D.M., C.W.V.K., M.S.G., A.P.S.).

## Abbreviations

AG: amylopectin granule
CDPK2: calcium dependent protein kinase2
COMP: TgLaforin-complemented parasite line
CRISPR: clustered regularly interspersed short palindromic repeats
DBA: *Dolichos biflorus* agglutinin lectin
ΔTgLaf: TgLaforin-KO parasite line
DHFR: dihydrofolate reductase
FACS: fluorescence activated cell sorting
GAA: acid-α-amyloglucosidase
Gal-NAc: N-acetylgalactosamine
GAPDH1: glyceraldehyde-3-phosphate dehydrogenase
GC/MS: gas chromatography/mass spectrometry
GFP: green fluorescent protein
GT1: glucose transporter1
GWD: glucan, water di-kinase
HA: hemagglutinin
HFF: human foreskin fibroblasts
HK: hexokinase
HR: homologous recombination
HXGPRT: hypoxanthine-xanthine-guanine phosphoribosyl transferase
IF: immunofluorescence
i.p.: intraperitoneally
KO: knockout
MPA: mycophenolic acid
PAM: protospacer adjacent motif
PAS: periodic acid-Schiff
PV: parasitophorous vacuole
PWD: phospho-glucan, water di-kinase
PYK1: pyruvate kinase1
RBC: red blood cell
RT: room temperature
SEX4: starch-excess4
SS: starch/glycogen synthase
TCA: tricarboxylic acid
TEM: transmission electron microscopy
UTR: untranslated region
WT: parental ME49ΔHXGPRT parasite line used in this study.

