## Supplementary figures and images for "TgLaforin, a glucan phosphatase, reveals the dynamic role of storage polysaccharides in *Toxoplasma gondii* tachyzoites and bradyzoites"

### Supplemental Figure 1

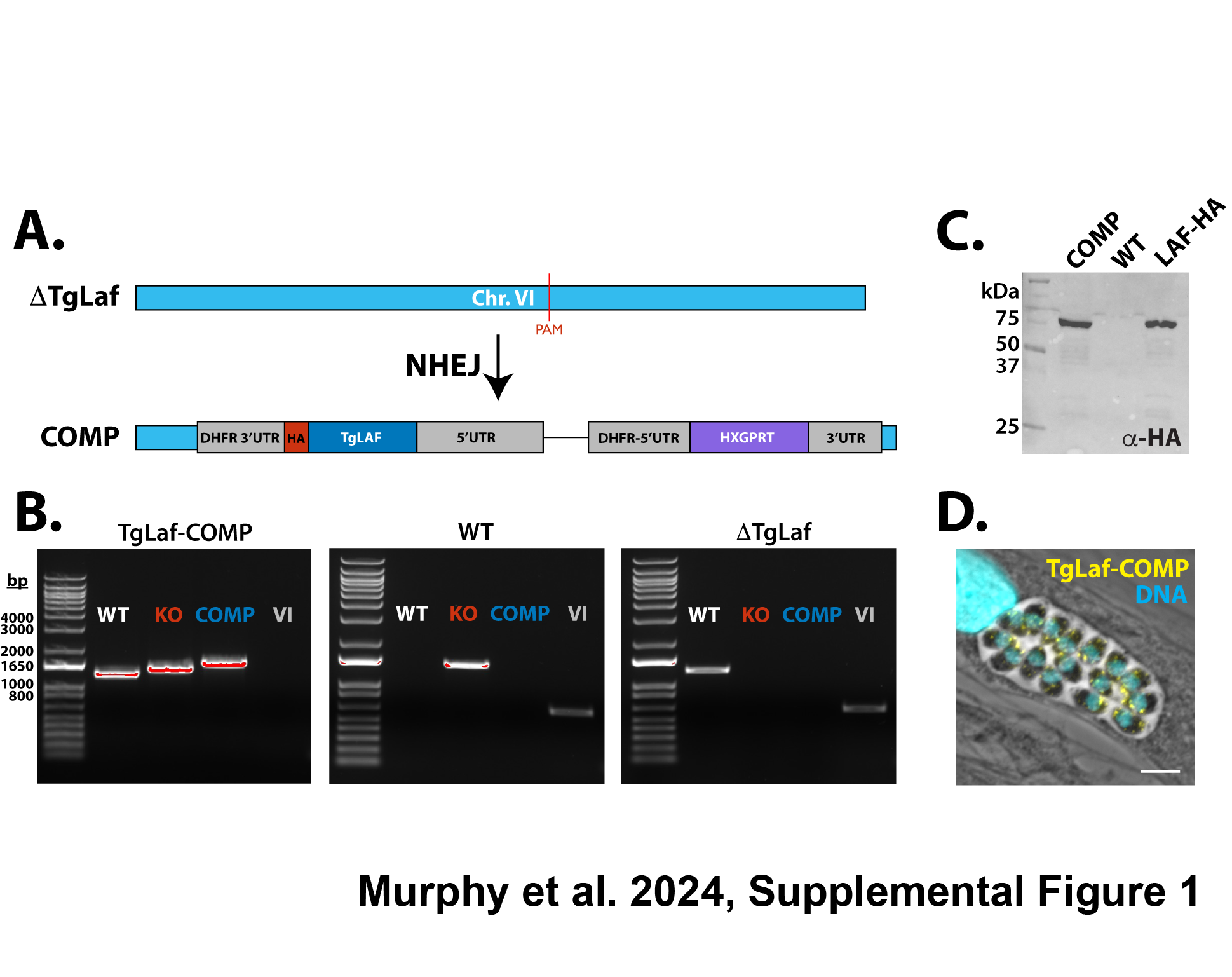

### Supplemental Figure 2

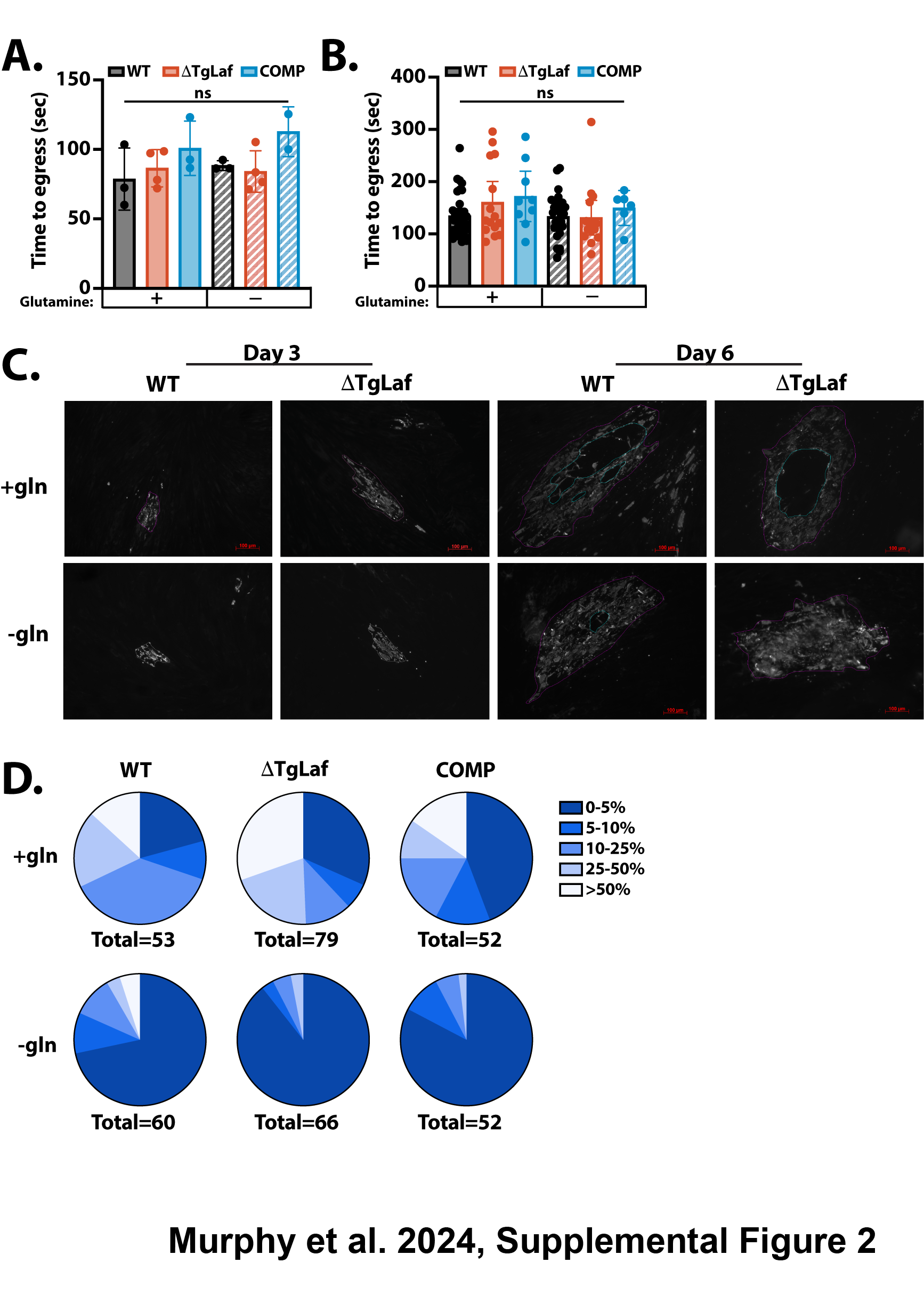

### Supplemental Figure 3

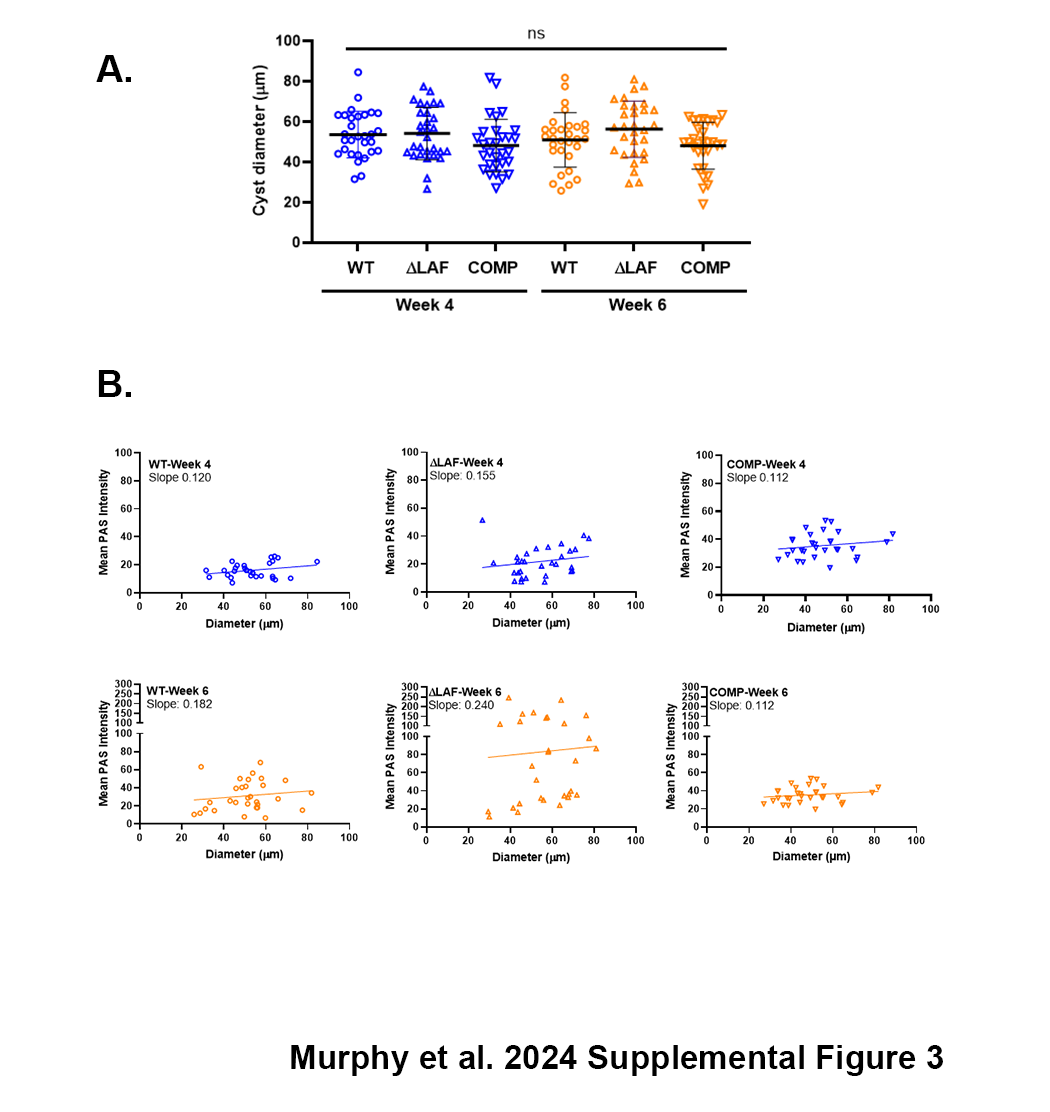

### Supplemental Figure 4

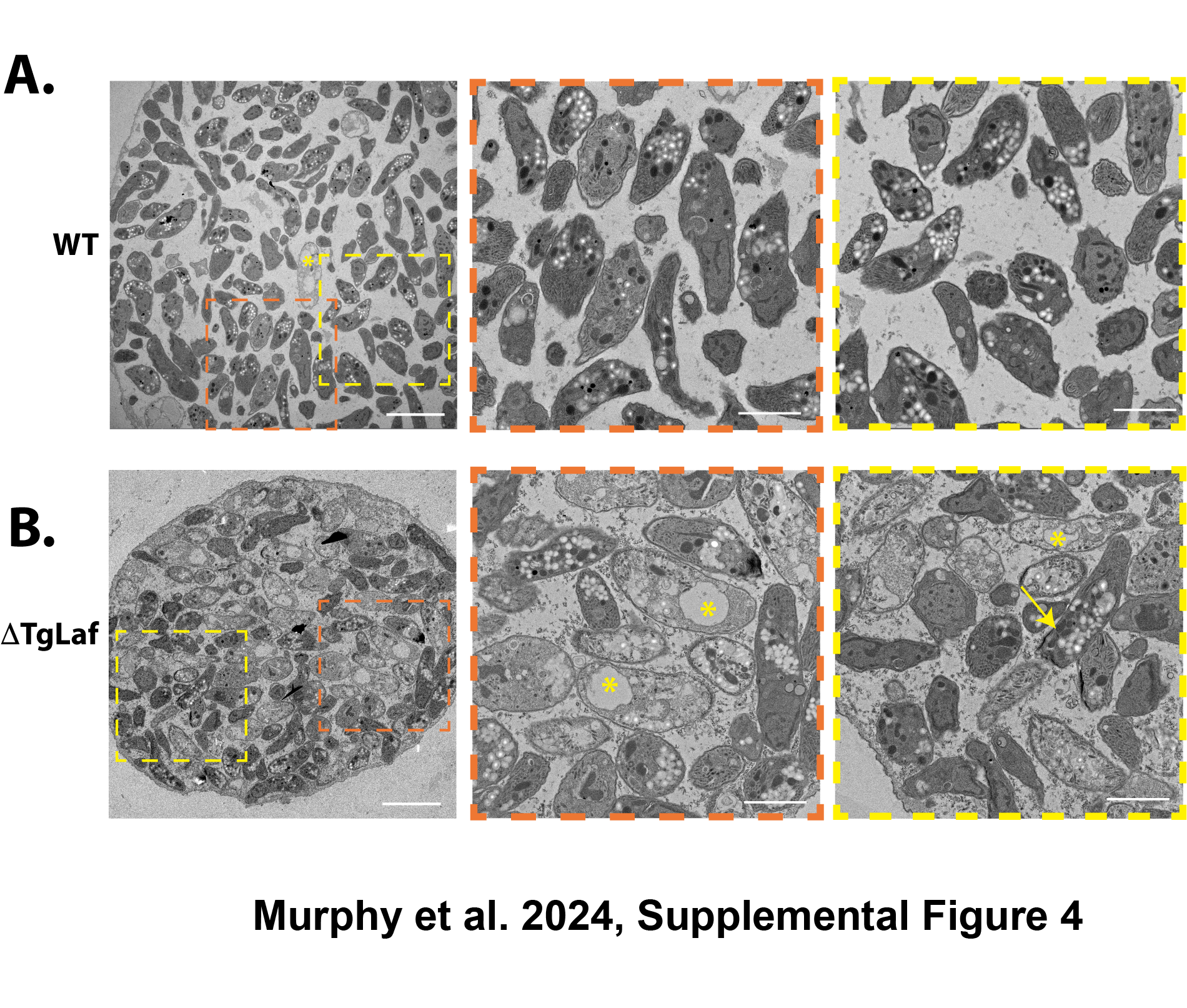
