## Supplemental File 5-Table 1 for "TgLaforin, a glucan phosphatase, reveals the dynamic role of storage polysaccharides in *Toxoplasma gondii* tachyzoites and bradyzoites"

**Supplemental Table 1. sgRNA sequences used in this study.**

| Target | Use | Sequence | Strand |
|---|---|---|---|
| TgLaforin 3'UTR | TgLaforin epitope tag | GCTTAGCGTGTGAACAGCAG | + |
| TgLaforin (exon 1) | TgLaforin KO | GAAGTCCCGATAACCTACGC | + |
| Chromosome VI | TgLaforin complementation | GCCGTTCTGTCTCACGATGC | + |
