## Supplemental File 6-Table 2 for "TgLaforin, a glucan phosphatase, reveals the dynamic role of storage polysaccharides in *Toxoplasma gondii* tachyzoites and bradyzoites"

**Supplemental Table 2. Plasmids used in this study.**

| Name | Use | Source |
|---|---|---|
| pSAG1::CAS9-U6::sgUPRT | Genetic modifications in *T. gondii* | David Sibley, Washington University |
| pJET-NcGra7_DHFR | TgLaforin knockout via DHFR-TS* knock in | Peter Bradley, UCLA |
| pHA3x-LIC | Tagging and complementation of TgLaforin | Peter Bradley, UCLA |
| TgLaforin-HA3x-LIC | Tagging and complementation, derived from pHA3x-LIC | GenScript |
