## Supplemental File 7-Table 3 for "TgLaforin, a glucan phosphatase, reveals the dynamic role of storage polysaccharides in *Toxoplasma gondii* tachyzoites and bradyzoites"

**Supplemental Table 3. Primers used in this study.**

Primers are presented in uppercase where the primer binds directly to the template, and in lowercase-bold where new sequence is being introduced. F, forward sequence; R, reverse sequence.

| Name | Use | Sequence (5' to 3') |
|---|---|---|
| TgLaf_3'UTR_sgRNA_F | sgRNA mutagenesis for TgLaf-HA epitope tagging | **tgaacagcag**GTTTTAGAGCTAGAAATAGC |
| TgLaf_3'UTR_sgRNA_R | | **cacgctaagc**AACTTGACATCCCCATTTAC |
| TgLaf_exon5_homology_F | Generation of TgLaf-HA tagging construct | AGAGGAGGCGGAGGAGAG |
| TgLaf_3'UTR_HX+homology_R | | TCCGTATCGCCCCCTCTCGTCTGACACGCCCTCTTTCCTCCAGCACGAAACCTTGCATTC |
| TgLaforin_sgRNA_E1_F | sgRNA for TgLaf-KO | **taacctacgc**GTTTTAGAGCTAGAAATAGC |
| TgLaforin_sgRNA_E1_R | | **tcgggacttc**AACTTGACATCCCCATTTAC |
| NcGra7-DHFR*_Fwd-TgLaf_homology | Amplification of NcGra7-DHFR* flanked with 40-nt TgLaforin | **gtctcttttctgcgcgtctccctgctcgtgcgtagaagg**CCACTCCATGGAACCTGACTG |
| NcGra7-DHFR*_Rev-TgLaf_homology | | **gctctttcccccattttcttttctctctcacgggttccg**CCTGCAAGTGCATAGAAGGAA |
| TgME49_ChrVI_sgRNA_F | sgRNA for ChrVI used in TgLaf complementation | **ctcacgatgc**GTTTTAGAGCTAGAAATAGC |
| TgME49_ChrVI_sgRNA_R | | **acagaacggc**AACTTGACATCCCCATTTAC |
| TgLAF_WT_F | "PCR1": Figure 3A. Amplifies WT TgLaforin | TCCTACATTCTGGAGCGAAG |
| TgLAF_WT_R | | AAAGCCACTTTCTCCAGGAG |
| DHFR*_R | "PCR2": Figure 3A. Amplifies ΔTgLaf chimeric locus in conjunction with TgLAF_WT_F. | GCATTATGAGGAAAGCCCAC |
| TgChrVI_WT_F | Amplification of WT Chr VI locus: ("VI": Figure | CAGGAAATATGCTGCGAGGA |
| TgChrVI_WT_R | | TGTGTCTGCTCTTGAAGGTG |
| TgLaf_COMP_F | Binds to HXGPRT cassette within complementation construct (derived from pHA3x_LICs | TGCAAGCCCTACATTGACAA |
